# Tool-use brain representations are independent of the acting body part and motor experience

**DOI:** 10.1101/2025.01.22.634305

**Authors:** Florencia Martinez-Addiego, Yuqi Liu, Kyungji Moon, Elizabeth Shytle, Lénia Amaral, Caroline O’Brien, Sriparna Sen, Maximilian Riesenhuber, Jody C. Culham, Ella Striem-Amit

## Abstract

The sensorimotor system is broadly organized somatotopically. However, an action-type organization has also been found: a division based on action-type independent of acting body parts has been shown for reaching and grasping actions. Does this generalization extend to non-ethological actions? Here, we examined fMRI responses for tool-use actions that participants performed with their hands or feet. We additionally tested individuals born without hands to control for hand motor imagery when performing foot actions. We show that the primary sensorimotor cortices have hand and foot selectivity, consistent with a somatotopic organization. In contrast, higher-level motor areas within the tool-use network, such as the premotor cortex, supplementary motor area, and superior parietal cortices, showed a shared preference for tool-use independent of the executing body part and sensorimotor experience. Multivariate decoding of action-type in these areas generalized between controls’ hand and foot and was successful in individuals born without hands. Finally, the temporal dynamics pattern in primary and association areas carried effector-specific and action-type information, respectively. Altogether, we show that the tool-use network in motor association areas represents higher-order action information beyond concrete motor parameters associated with specific effectors, and regardless of hand motor experience. This suggests that an action-type, effector-independent organization extends beyond ethological actions, supporting a hierarchical organization in the action domain. Further, it shows that functional organization in congenital handlessness is based on the hierarchical organization of the intact cortex.

## Introduction

During action planning, our motor system effortlessly translates conscious goals to elaborate, coordinated muscle contractions. There is evidence suggesting that a hierarchical organization in the motor system facilitates this translation (Poggio & Bizzi, 2004; Grafton & Hamilton, 2007; Friston, 2008; Giszter, 2015; Merel et al., 2019): basic motor features are encoded within the primary motor cortex (M1) and higher-level association areas represent more complex information such as the action-type (Evarts et al., 1968; Hatsopolous et al., 2007; Merel et al., 2019). M1 has a broadly somatotopic organization, whereby each body part is represented by a distinct area of the cortex, and adjacent body parts are mapped to neighboring cortical regions (Penfield et al., 1937; but see Gordon et al., 2023). This somatotopic organization in M1 represents lower-level, body-part-specific kinematic information by encoding muscle synergy information (Holdefer & Miller, 2002; Overduin et al., 2015; Leo et al., 2016; Gallego et al., 2017).

Motor association areas, such as the dorsal premotor cortex (PMd), ventral premotor cortex (PMv), supplementary motor area (SMA), and intraparietal lobule (IPL), instead represent goal-directed action information such as target location (Alexander & Crutcher, 1990; Kakei, Hoffman, & Strick, 2001; Kakei, Hoffman, & Strick, 2003; Cisek et al., 2003), motor sequence (Mushiake, Inase, & Tanji, 1990), and action type (Rizzolatti et al., 1998). As expected from areas higher in the motor hierarchy, there is evidence that these representations are partly invariant to the low-level motor features. In recent years, there has been accumulating evidence that these areas may contain action representations that generalize across executing body parts (effectors). During motor planning, the response patterns in the posterior parietal cortex (PPC) and PMd encode the upcoming action-type (e.g., reaching or grasping) regardless of which hand would be used (Gallivan et al., 2013a). For the execution of reaching and grasping actions, the premotor cortex and parietal lobe encode target location, target size, and orientation independent of whether an action was executed by the dominant or non-dominant hand (Haar et al., 2017; Turella et al., 2020). Additionally, the PMd and intraparietal sulcus (IPS) are involved in the planning of both hand and eye movements towards a specified direction (Gallivan et al., 2011; Heed et al., 2011; Leoné et. al., 2014; Gallivan et al., 2016; Heed et al., 2016; Magri et al., 2019). Importantly, these findings were not limited to effectors that are somatotopically close, homologous, or frequently involved in coordinated actions (i.e., hand-hand and hand-eye coordination). A few studies have found common activation in frontoparietal areas during action planning for pointing movements between the hand and foot, which are somatotopically distant and less coordinated (Heed et al., 2011; Heed et al., 2016; Leoné et al., 2014). A shared action representation across the hand and foot has been extended to include the differentiation of reaching and grasping action-types. We have recently shown that foot reaching and grasping actions performed by individuals born without hands engaged some of the same areas as hand actions performed by controls (Liu et al., 2020). Together, these findings suggest that motor association areas support reaching and grasping actions independently of the performing body part.

These results provide a theoretical grounding for a functional, action-type organization of motor association areas. In this scenario, the action-type (e.g., reaching) and not the acting effector would be the leading organizational principle (Kaas, 2013; Graziano, 2016; Hahamy et al., 2017; Liu et al., 2020). Importantly, evidence supporting effector-independent action-type representations has so far been limited to these two specific ethological, evolutionarily driven actions that are shared across non-human primates (Kaas, 2013; Graziano, 2016). Yet, humans have developed additional motor skills, such as sophisticated tool-use, with dedicated neural substrates (Johnson-Frey, 2004; Orban and Caruana, 2014; Peeters et al., 2009). It is unclear whether motor association areas will also support an effector-independent action-type representation for additional actions, including this newly evolved action-type.

Generally, tools are defined as instruments that facilitate the completion of goals (for a deeper discussion on definitions of tools, see Osiurak et al., 2010). Tool-use marks an important evolutionary milestone for humans (Ambrose, 2001). Indeed, though tool-use ability is shared with non-human primates (Goodall, 1964; Seed & Byrne, 2010), humans have developed a unique dexterity, expertise, and understanding of the causal relationship between tool-use and the end outcome (Povinelli et al., 2000). This is reflected neurally as there are different neural substrates for tool-use in humans versus non-human primates (Peeters et al., 2009). Humans have an expanded network of parieto-frontal areas (Krubitzer and Prescott, 2018), including the supramarginal gyrus, middle temporal gyrus, aIPS, and PMv that is engaged in tool-use (Ishibashi et al., 2016; Reynaud et al., 2016; Gallivan et al., 2013b; Lewis, 2006).

Here, we test the hypothesis that the tool-use network is also organized by action-type, regardless of the executing effector. Evidence supporting this hypothesis would suggest that an action-type, effector-independent organization is not limited to ethological actions and further argue for a motor abstraction hierarchy. Alternatively, generalizing across effectors for the same action may only be possible for reaching and grasping, or even more broadly just for ethological actions. This instead would suggest longer evolutionary timelines are needed for action-type specialization.

In this study, we used functional neuroimaging to investigate whether motor association areas support effector-independent tool-use representations. To this end, we recruited a group of typically developed individuals who performed in-scanner tool-use and basic grasping actions with their hands (primary effector) and their feet (atypical effector). Though a shared preference for tool-use between the hands and feet in typically developed individuals would provide evidence for effector-independent brain mechanisms, there is an alternative explanation. When asked to perform typical hand actions with the foot, the participant may imagine executing the action with their hand and only then translate it to the foot, resulting in shared activation with hand execution that is a product of hand motor imagery (Ehrsson et al., 2003; Lacourse et al., 2005). Likewise, any differential, effector-dependent activations may stem from a difference in experience between the hand and the foot. To avoid these possible confounds, we leveraged a population of individuals born without arms and hands (individuals with upper-limb dysplasia, abbreviated IDs) who use their feet as their primary effector. Since the IDs cannot imagine moving their own hands, including them allows us to bolster claims about effector-independent and effector-dependent networks. By mapping effector-independence and dependence we can test if the motor hierarchy generalizes across body parts for non-ethological, human-specific actions.

## Results

To address our main question of whether the brain supports tool-use action-type representations that generalize across body parts (effectors), we scanned individuals with dysplasia (IDs, born without hands; n = 6) and typically developed control participants (n = 22, data excluded for five; see **Methods**). Participants executed grasping and simple tool-use actions (turning a spatula, see schematic in **Fig. S1**). The recruited IDs use their feet as the compensatory body part to perform daily actions (Vannuscorps et al., 2014; Striem-Amit et al., 2017; Striem-Amit et al., 2018; Vannuscorps & Caramazza, 2019; Vannuscorps et al., 2019; Liu et al., 2020) and therefore completed all in-scanner actions with their dominant foot (five IDs were right-footed, one ID was left-footed). Control participants were right-handed and right-footed and completed actions using their right hand or right foot in separate experimental runs. We used several approaches to test effector-independence in tool-use actions: (1) univariate main effects of effector and action-type, and how these properties are shared across the groups; (2) consistent preference for tool-use across controls’ hand, controls’ foot, and IDs’ foot data; (3) multivariate decoding of the action-type within and across effectors; (4) shared neural temporal signals for tool-use regions across effectors; and (5) decoding of action-type from the temporal dynamics pattern across effectors. Together, these analyses provided converging results supporting the existence of effector-independent action representations for tool-use.

## Univariate effects of effector and action-type

We performed two random-effects (RFX) ANOVAs to determine the brain regions showing 1) a preference for a specific effector and 2) selectivity for action-type. One ANOVA was computed for control participants’ data with effector (hand and foot) and action-type (grasping and tool-use) as within-subject factors (CH+CF; **Fig. 1A)**. Since typically developed individuals are not accustomed to using their feet for grasping and tool-use actions, they might engage hand imagery, and hence hand areas, to perform their foot actions. In this case, a finding of effector-independence would be less interpretable. To control for this potential confound, an additional ANOVA was performed across the controls’ hand data and IDs’ foot data with effector/group as a between-subjects factor and action-type as a within-subjects factor (CH+IDF; **Fig. 1D)**. For each main effect, we calculated the spatial overlap between the two ANOVAs to investigate effector-dependence and effector-independence that generalizes across experience and dexterity while ruling out potential manual motor imagery.

**Figure 1.**
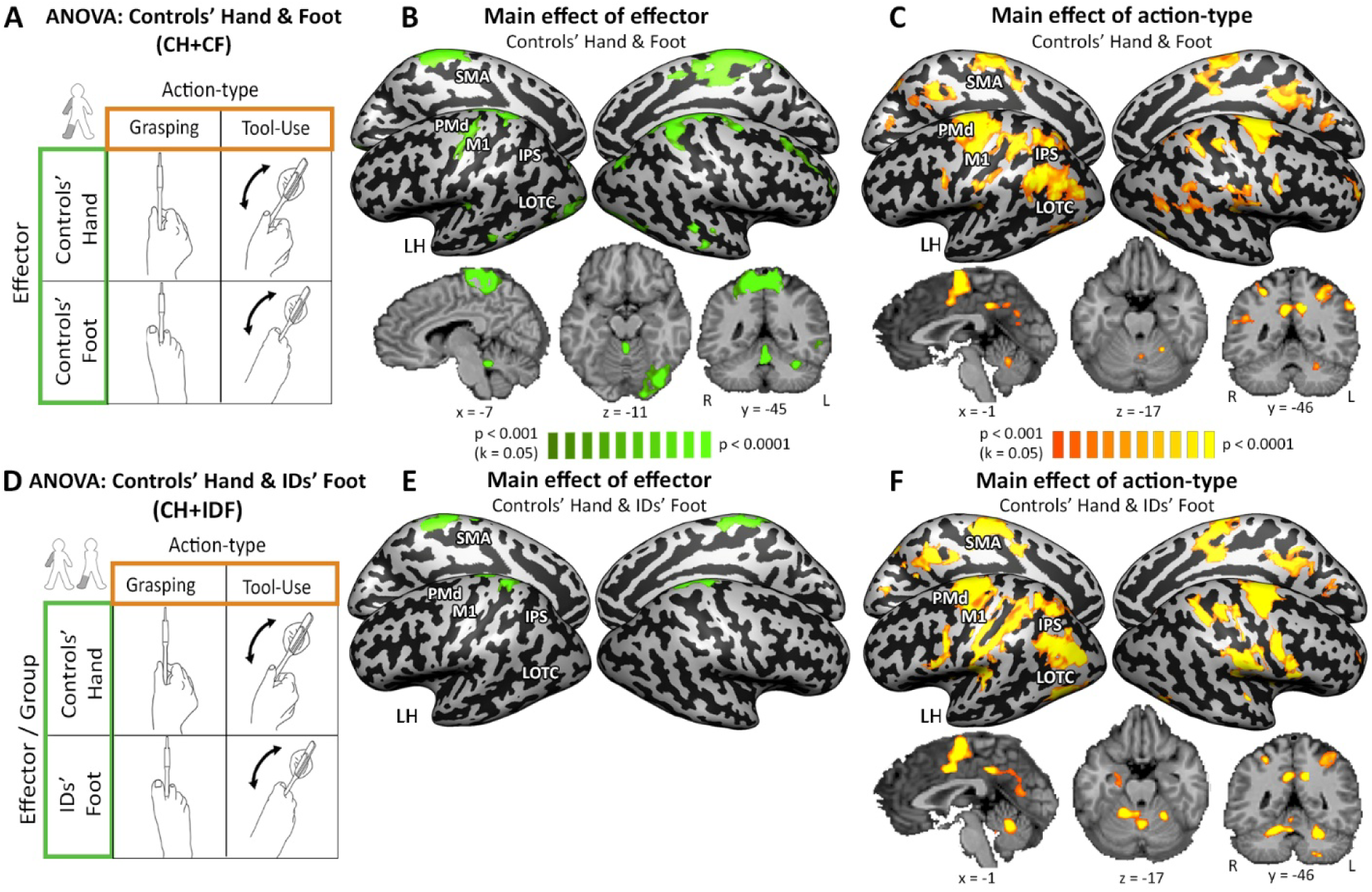
Effector-dependent and effector-independent action networks for tool-use and grasping. Results from RFX ANOVA analyses for controls’ hand and controls’ foot (A) as well as controls’ hand and IDs’ foot (D). Each map was thresholded at *p* < 0.001 and corrected for multiple comparisons at *p* < 0.05. Two RFX ANOVAs were performed. (A) One ANOVA was within-group and consisted of controls’ hand and controls’ foot data. In this ANOVA, action-type (tool-use, grasping) and effector (hand, foot) were within-subjects’ factors. (D) The second ANOVA was across groups and consisted of controls’ hand and IDs’ foot data. In this ANOVA, the effector was a between-subjects factor and action-type was a within-subjects factor. (B and E) Main effect of effector. (B) Difference between hand and foot actions in controls was found in left M1 (foot and hand area), SPL, and LOTC as well as left cerebellar areas IV and VI. Data analyses were conducted at the volume level and projected to cortical surfaces for visualization purposes. (E) Difference between hand actions in controls and foot actions in IDs (right) was found left M1 (foot area) and the SPL. (C) A shared preference for action-type independent of executing effectors across controls’ hand and foot was found in bilateral PMd, IPS, SMA, and LOTC as well as areas left V and left VI of the cerebellum. (F) A shared preference for action-type independent of executing effector (hand for controls, foot for IDs) and sensorimotor experience was found in bilateral PMd, IPS, SMA, and LOTC as well as areas left V, VI, and VIIIb of the cerebellum.

We first probed each of the ANOVAs for areas with a main effect of effector across actions (consistent activation preference based on the acting body part), which indicated an effector-dependent representation. In the CH+CF ANOVA, we found a differential response based on the acting body part in the left primary sensorimotor (M1/S1 hand and foot regions), left lateral occipital temporal cortex (LOTC), left superior parietal lobule (SPL), right SMA, and lobules IV and VI in the left cerebellum (**Fig. 1B**). Similarly, the CH+IDF ANOVA showed differential activations based on the acting effector in the left foot primary sensorimotor cortex and the left SPL (**Fig. 1E**). Accordingly, the two ANOVAs overlapped in the left foot primary sensorimotor area the left SPL **(Fig. 2A)**, suggesting an effector-dependent organization in these regions. Although at the whole-brain level there was no main effect of effector in hand-selective M1 at the chosen statistical threshold, it was found when directly sampling the hand M1 region (F(1,21)=9.89, *p<0.005*).

**Figure 2.**
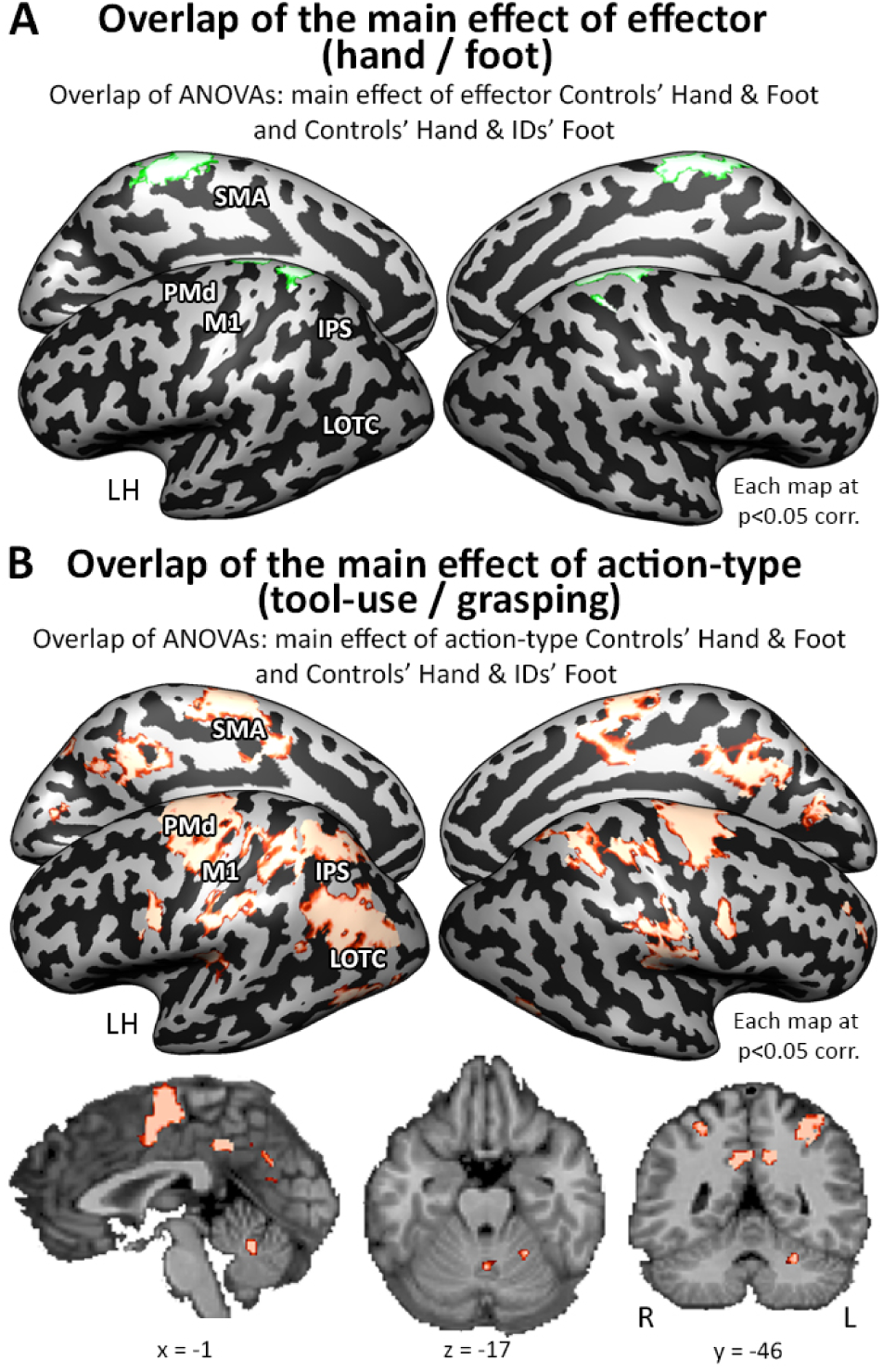
Consistent Networks Across Effectors and Sensorimotor Experience. Shared results from RFX ANOVA analyses for controls’ hand and controls’ foot as well as controls’ hand and IDs’ foot. (A) Overlap of main effects of effector from Fig. 1B and 1E. A consistent effector effect is found in the left M1 (foot area, as well as hand area; see results) and the SPL (area 5L). (B) Overlap of main effects of effector from Fig. 1C and 1F. A consistent action effect is found in bilateral PMd, IPS, SMA, and left LOTC as well as lobules V and VI of the left cerebellum.

We then asked if there was evidence for a functional, effector-independent organization for tool use. To this end, we examined brain areas showing a main effect of action-type regardless of the executing effector (consistent activation preference based on the performed action). Results from the CH+CF ANOVA revealed a widespread frontoparietal network containing the bilateral SMA, PMd, SPL and LOTC, as well as lobules V and VI of the left cerebellum **(Fig. 1C)**. The CH+IDF ANOVA showed a similar pattern of widespread frontoparietal network including the bilateral IPS, PMd, and SMA and the left LOTC, as well as lobules V, VI, and VIIIb in the left cerebellum **(Fig. 1F)**. Overlapping the two ANOVAs, we found that the bilateral SMA, PMd, SPL, and left LOTC along with lobules V and VI of the left cerebellum showed selectivity to action-type regardless of the effector and sensorimotor experience **(Fig. 2B)**. Importantly, these effects were not simply driven by a potentially longer movement duration in tool use vs. grasping, as consistent results were found in a control analysis accounting for the movement onset and duration (**Fig. S2**). Notably, these areas do not show a significant interaction effect (**Fig. S3**), further corroborating effector-independence. We also computed a supplementary RFX ANOVA comparing the controls’ foot and IDs’ foot actions (**Fig. S4**) and found a consistent main effect of action-type independent of limb experience in all the same areas. Several of these regions, namely the PMd, SPL, and anterior SMG, are known to be part of the tool-use network (Lewis, 2006), suggesting that tool-use actions are represented by this network at a functional, effector-independent level.

## Consistent tool-use preference across effectors

Though above regions have been shown to be part of the tool-use network, the ANOVA main effects analysis does not inherently specify the direction of action preference. As a result, it is unclear which action is driving the observed main effect. To directly investigate whether tool-use areas may be effector-independent, we used controls’ hand data to define tool-use selective regions-of-interest (ROIs) and tested for a consistent preference for tool-use over grasping in foot data. ROIs that supported tool-use in controls’ hand data included core tool-use areas in the left hemisphere – such as SMA, PMd, SPL (7pc), SPL (area 2), PMv, and LOTC (see **Fig. 3A**, **Table S1**; for additional areas that showed a tool-use preference, see **Fig. S5**, **Table S2**, ROI labels were drawn from the Julich Brain Atlas at the volume level; Amunts et al., 2020). Within these areas, we found a consistently higher response for tool-use over grasping for both controls and IDs even though they were executing the actions with their feet (**Fig. 3B**; for detailed statistics see **Table S3**). Notably, the findings from PMd, SPL (7pc), and SPL (area 2) are robust to the correction of recorded movement onset and duration (**Fig. S6A**).

**Figure 3.**
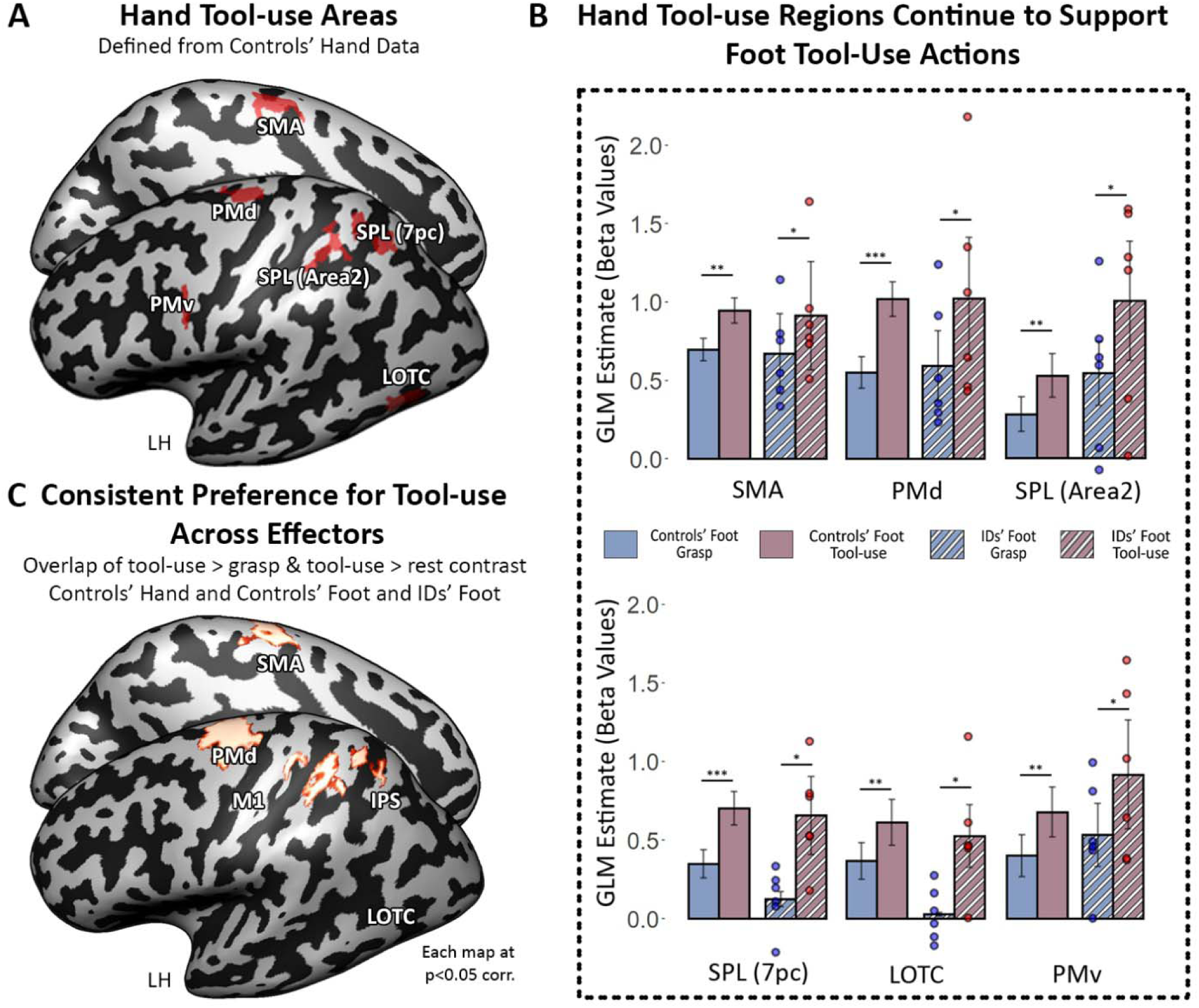
Tool-use Areas Have Effector-Independent Selectivity. (A) Tool-use ROIs defined from CH data that show a main effect of action-type and have stronger activity for tool-use over grasping (map was thresholded at *p* < 0.01 and corrected for multiple comparisons at *p* < 0.05). (B) Bar graphs of average GLM beta estimate values for the tool-use ROIs shown in (A) for controls’ foot (solid bars) and IDs (striped bars). Each dot above a striped bar plot represents one individual with dysplasia. Black asterisks are FDR-corrected. *: *p* < 0.05; **: *p* < 0.01; ***: *p* < 0.001. (C) A three-way spatial overlap of a direct contrast of tool-use > grasp (in conjunction with tool-use vs. rest) is shown across controls’ hand, controls’ foot, and IDs’ foot group data (each at p < 0.05 corr.). A consistent preference for tool-use was found in the left PMd, SMA, and SPL.

To explore if additional areas show effector-independent tool-use preference, we performed a direct contrast of tool-use > grasp (in conjunction with tool-use > baseline to include only positive activations) for controls’ hand, controls’ foot, and IDs’ foot data separately, and computed a three-way overlap among them. We found a consistent, significant preference for tool-use across effectors in the left PMd, SMA, and SPL (areas 2 and 7pc) (**Fig. 3C**; for each group’s contrast see **Fig. S7**). This result was replicated when controlling for action duration (**Fig. S6B**). Out of the six IDs, significant tool-use preference at the single-subject level was found in all, five, and four individuals in the left PMd, left SMA and SPL, and in LOTC, respectively (**Fig. S8**). Together, this shows that canonical hand tool-use regions can generalize to other body parts.

Across analyses, these findings suggest that tool-use areas, namely PMd, SPL (7pc), and SPL (area 2), generalize across the acting body parts and have a shared preference for tool-use across effectors.

## Decoding action-type across effectors

Next, we leveraged multi-voxel pattern analysis (MVPA) to determine if there are consistent, distinguishable, spatial neural response patterns for the different actions that extend across acting body parts. We first performed ROI-MVPA in the hand tool-use selective regions defined in **Fig. 3A** (**Fig. 4A**). To do this, we trained a linear support-vector machine (SVM) classifier to differentiate between grasping and tool-use either within an effector (controls’ hand, controls’ foot, IDs’ foot), or across effectors in controls (SVM was trained on hand data and then tested on foot data and vice versa, see **methods**). Given the possibility for additional noise from motion in the data that the classifier may be sensitive to, thereby artificially increasing decoding, we further baselined the accuracies to a control ROI, the white matter of the bilateral ATL. As our task engages widespread visual, auditory, and motor networks, the bilateral ATL is among the few brain areas that are not expected to respond to our paradigm. For baselining, we subtracted each participant’s decoding accuracy in ATL from that of each ROI, thereby maintaining within-subject variance. We accordingly tested for significantly greater decoding accuracy in the ROIs compared to ATL and chance level. In all hand tool-use selective regions, action-type decoding was successful within controls’ hand and foot, and across effectors. Importantly, the IDs also had significant action-type decoding in the PMd, SPL area 7pc, and SPL area 2 (**Fig. 4A**).

**Figure 4.**
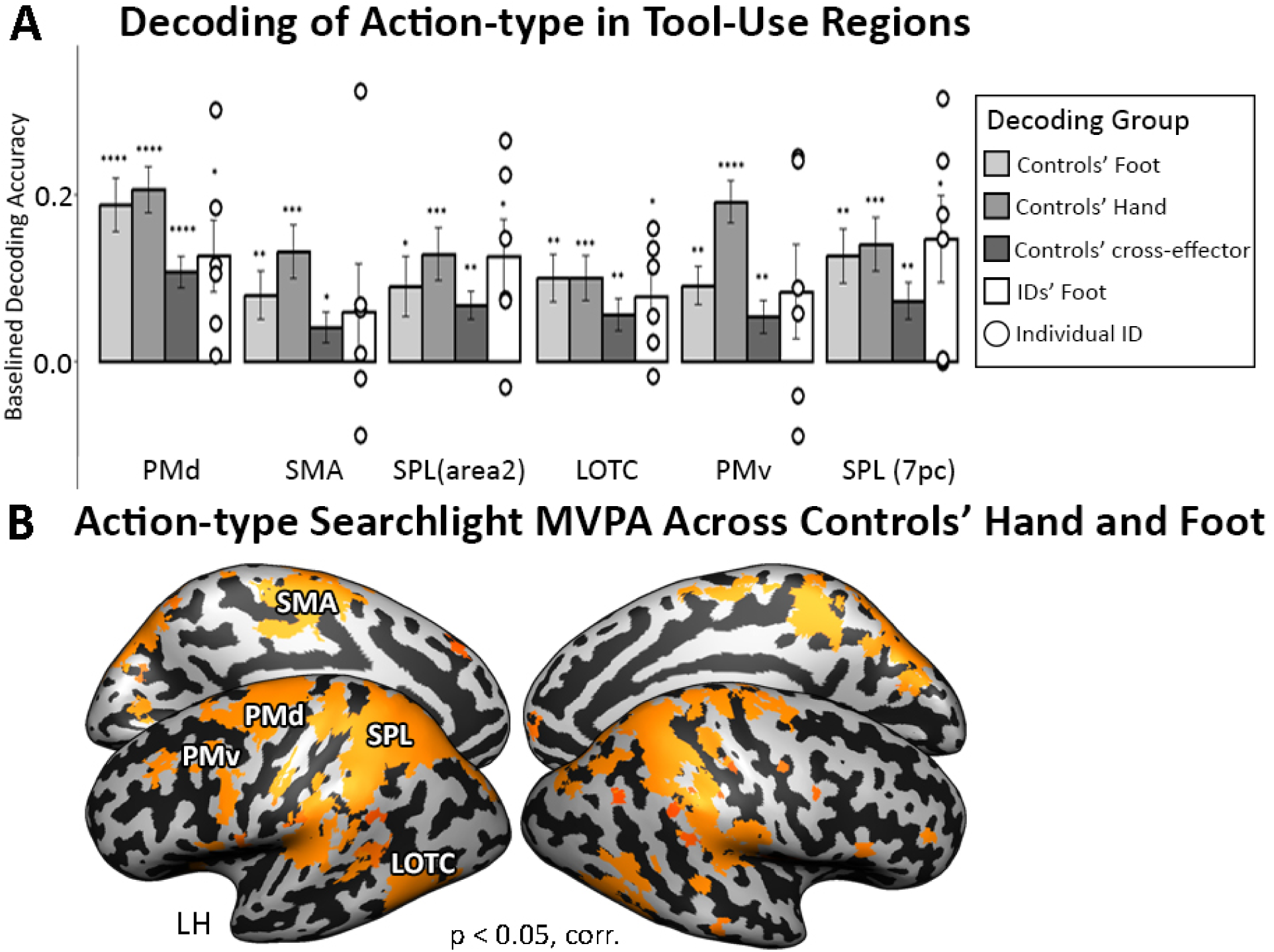
Multi-voxel pattern analysis reveals action code shared between the hand and the foot. (A) Tool-use regions show significant action decoding regardless of effector. Decoding of action-type (tool-use vs grasp) in controls (within hand and within foot data) and IDs (within foot data) and across effectors (across controls’ hand and foot) is significant in tool-use regions as identified by univariate analyses (same regions as used in Fig. 3A,B). Accuracies were baselined to a control ROI (the bilateral ATL) and to chance level (50%) Black asterisks are FDR-corrected. *: *p* < 0.05; **: *p* < 0.01; ***: *p* < 0.001. (B) Areas with significant cross-effector decoding of action-type in the control group (as compared to chance). Error bars denote SEM.

Further, we performed a whole-brain searchlight MVPA decoding action-type across effectors in the control group, as compared to chance (50%). This confirmed the decoding in the tool-selective network and additionally revealed a shared neural code for action-type independent of the executing effector in the bilateral inferior parietal lobe and PMv (**Fig. 4B**). Together, these results robustly show effector-independent neural patterns distinguishing the two actions, regardless of the body part or sensorimotor experience.

## Differential time courses across the action hierarchy

Going beyond typical univariate and multivariate analyses, we tested whether evidence for an action hierarchy, namely effector-dependent representations in primary motor areas and effector-independent representations in motor association areas, would also manifest in temporal dynamics of the fMRI response. Although BOLD signals are known for their low temporal resolution, a slow event-related design with a relatively short TR (782ms) in our experiment allowed us to detect differential time courses between the conditions.

If the association motor areas encode effector-independent action-type information, the timing of their involvement during the action stages should be similar, regardless of the acting effector. Given that our experimental design was structured such that all participants used the same trial order (for both hand and foot runs), we tested this hypothesis by using the average time course from an effector-independent region from controls’ hand data (PMd) as a predictor of the foot neural response in IDs in a whole-brain GLM analysis. The motor association cortices’ response, including PMd, PMv, and SPL, were synchronized to the same action stages for IDs’ foot actions as with controls’ hand (**Fig. 5A**). In contrast, the time-course in M1 hand area for controls’ hand predicted IDs’ foot responses along primary sensorimotor cortex, especially in foot M1 (**Fig. 5B**). These results suggest that higher and lower areas in the motor hierarchy share similar temporal dynamics in our task, regardless of being performed by the hand in controls or by the foot in IDs. A direct contrast between the two, showing differential timed response for the low-vs. high-level areas of the motor hierarchy, revealed a frontoparietal network in IDs where the time-course resembles PMd in controls’ hand and primary sensorimotor areas where the time-course resembles M1 in controls’ hand (**Fig. 5C**). These results show that the motor hierarchy responds with similar temporal dynamics across acting body parts.

**Figure 5.**
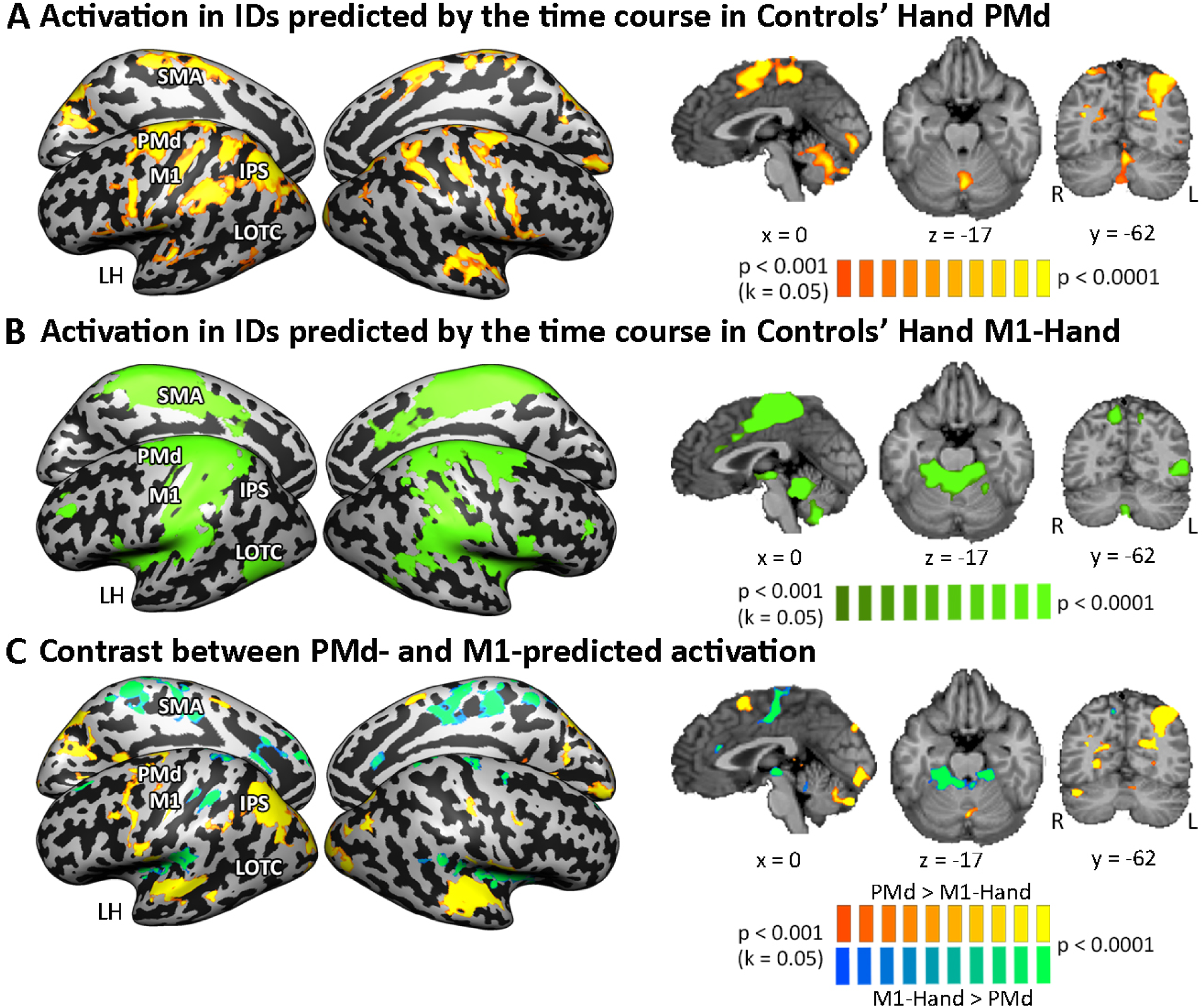
Similar temporal dynamics for motor regions across effectors. Time-courses from controls’ hand PMd and hand M1 correlate with parallel networks in IDs across the tasks. All maps were thresholded at *p* < 0.001 and corrected for multiple comparisons at *p* < 0.05. (A) Using a seed region of PMd (defined from tool-use preference; Fig. 3A), the time-course of controls’ hand data was used as a predictor for individuals with dysplasia using a GLM analysis, revealing a motor association network engaged at the same times during action dynamics. (B) Using a seed region of hand M1 (defined by body part preference from the CH+CF ANOVA in Fig. 1B), the time-course of controls’ hand data was used as a predictor for individuals with dysplasia, revealing the involvement of the low-level motor network at the same times during action dynamics. (C) A contrast of the two seeds’ predictors (PMd and M1) was performed to highlight differences between the two networks, highlighting similar temporal dynamics across the execution of the task with the hand in controls and the foot in IDs. See Fig. S10 for this analysis repeated with controls’ hand data.

## Temporal dynamics patterns along the action hierarchy

We further performed a temporal multivariate pattern analysis (tMVPA, Vizioli et al., 2018) to test if the temporal dynamics of the response pattern over the course of a trial carries effector or action-specific information. Tool-use action-selective ROIs (**Fig. 3**) and body-part selective areas (M1 hand and foot areas) defined from the controls’ univariate results (see **Methods**) were analyzed. Within each ROI and for each effector and action-type, we calculated a single-trial representational dissimilarity matrix (stRDM, see **Methods**) that depicts the distance (in 1 – Pearson’s correlation) in the multi-voxel response pattern between each pair of time points over the course of a trial. Trials were defined from the onset of the action instruction (e.g., “grasp”), through the planning period, the execution of the action, and the rest period following each trial. The pattern of the stRDM reflects the temporal dynamics: low dissimilarity along the diagonal indicates dynamic representations changing from time to time, and low dissimilarity also off the diagonal denotes stable representations over time. We then performed MVPA and hierarchical clustering on the stRDMs to test whether different effectors and action-types are associated with distinct temporal dynamics patterns.

We first tested the expectation that the primary motor cortex would show effector-specific temporal dynamics (**Fig. 6**). First, a two-phase temporal dynamics pattern, dividing each trial to two main portions and likely corresponding to planning and execution stages of the action, was observed only for the “native effectors”, i.e., hand actions in M1 hand area and foot actions in M1 foot area (stRDMs in **Fig. 6A, 7D**). Second, the action-type could be decoded based on the stRDMs in the M1 hand area mainly for controls’ hand (bar graph in **Fig. 6B**), and hand actions form distinct clusters away from the less distinguishable foot actions (hierarchical clustering in **Fig. 6C**). Importantly, effector/group information could be successfully decoded both within and across action-types (**Fig. S10A**), suggesting a general effector representation regardless of action. Interestingly, the M1 foot area also shows distinct dynamics patterns for the hand and a tendency of cross-group action decoding (bar graph in **Fig. 6E**). Nevertheless, decoding of effector/group was significantly above chance both within and across actions (**Fig. S10A**) in M1 foot area, and the hierarchical clustering shows a separate cluster for each effector, consistent with an effector-based temporal pattern (**Fig. 6F**, right panel). Finally, an omnibus hierarchical clustering analysis on the dissimilarity patterns of all three groups and two regions revealed a somatotopic organization such that the temporal dynamic patterns of the native effectors are clustered together within which the foot actions forming sub-clusters, and the non-native effectors form a separate cluster that is less differentiable and without a clear structure (**Fig. 6G**).

**Figure. 6.**
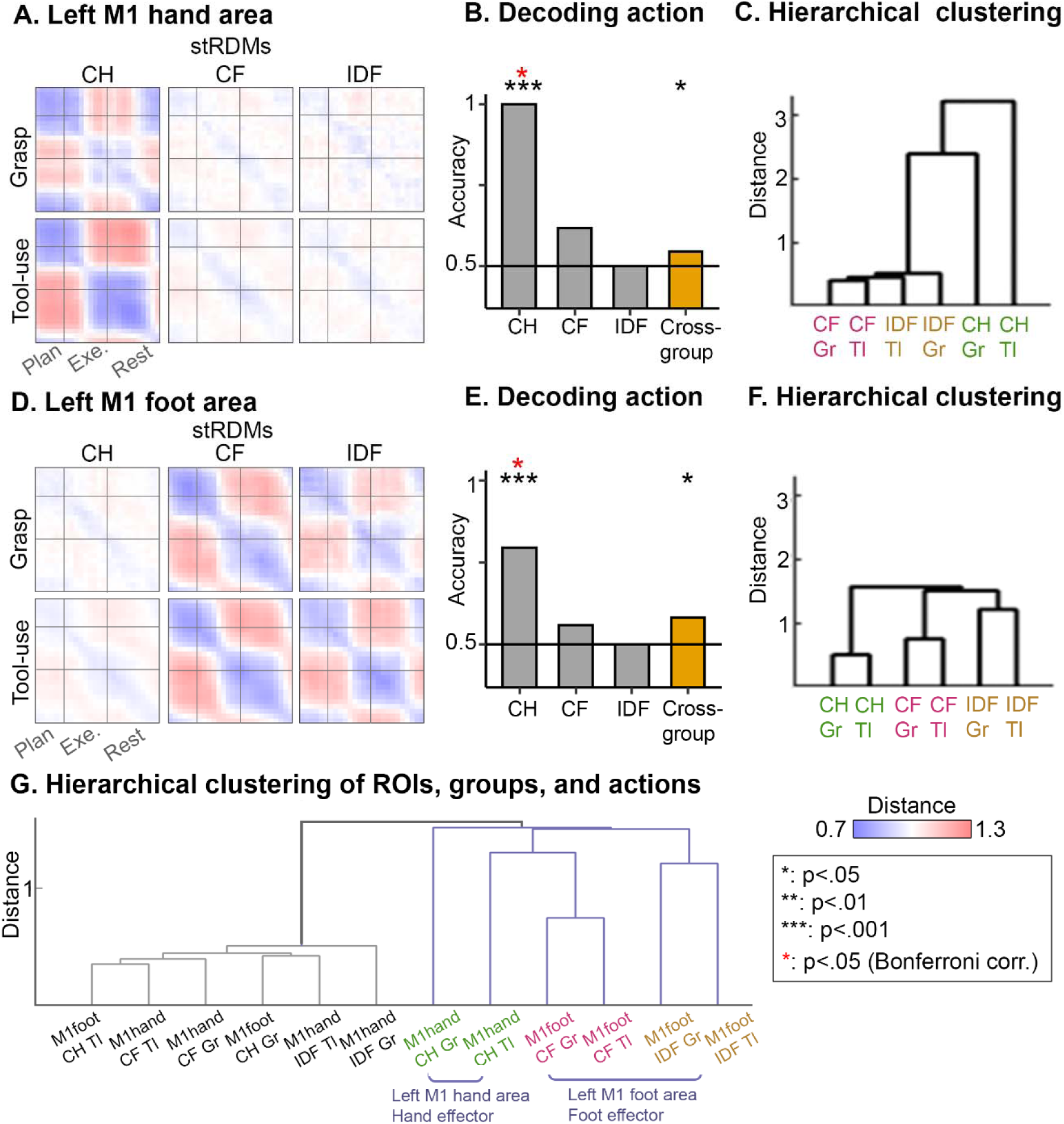
Temporal dynamics patterns in effector-dependent primary motor cortex. A-B: Data is shown for the hand (A-C) and foot (D-F) primary sensorimotor cortex. A and D: For each action-type and effector, a single-trial representational dissimilarity matrix (stRDM) is computed where the values are 1-Pearson’s r between the multivoxel response patterns of each pair of time points within a trial (left-most dissimilarity pattern). Different stages of the trials can be seen reflected in the dynamics, corresponding to a change between action planning and execution, as well as the end of execution and the beginning of the rest period after each trial. B and E: The bar graph depicts decoding accuracy of action-type within each effector and across all three groups/effectors (CH, CF, IDF) based on stRDM. C and F: Hierarchical clustering depicts the grouping of stRDMs based on their similarity, with longer lines along y-axis denoting larger dissimilarity between items/clusters. The M1 hand and foot area largely show effector-based decoding and hierarchical clustering. G: Hierarchical clustering performed on all stRDMs from M1 hand and M1 foot area. Two large clusters are formed, one grouping native effectors (right side, blue lines) and the other non-native effectors (left side, black lines). Within the native-effector cluster, each foot effector (CF and IDF) form a sub-group.

Importantly, the association motor areas display a different, action-based, temporal dynamics’ patterns across effectors/groups. Despite variable execution manners across effectors, grasping and tool-use stRDMs could be decoded within and across effectors in left PMd, IPS and SPL, providing evidence for a general action-type representation in the temporal domain (**Fig. 7B, E, H**). In contrast, effector/group information could no longer be decoded in these areas (**Fig. S10A**), forming a double dissociation between the primary and association motor areas in terms of effector-dependent and action-dependent temporal patterns. Finally, hierarchical clustering analyses revealed a tendency of grouping the temporal dynamics patterns based on the performed action-type as opposed to effectors (**Fig. 7C, F, I**). These results are unlikely simply driven by different execution times between actions and effectors, as we found a double-dissociation across brain regions. Further, to conclusively verify that this result could not have been driven by head motion induced by action execution, we performed a control analysis where we replicated the tMVPA analysis with stRDMs created using the six head motion parameters as the “response pattern”. This control analysis found a unique temporal pattern for head motion that is instantaneous to execution (**Fig. S10B**), and dissociable from the neural stRDMs. Moreover, action-type information could *not* be decoded in this control analysis, effectively ruling out the possibility that the temporal patterns simply reflect head movements. Altogether, these results revealed effector-dependent and effector-independent temporal dynamics patterns in the primary and association motor cortex, respectively, lending further evidence for a representational action hierarchy from the temporal domain.

**Figure 7.**
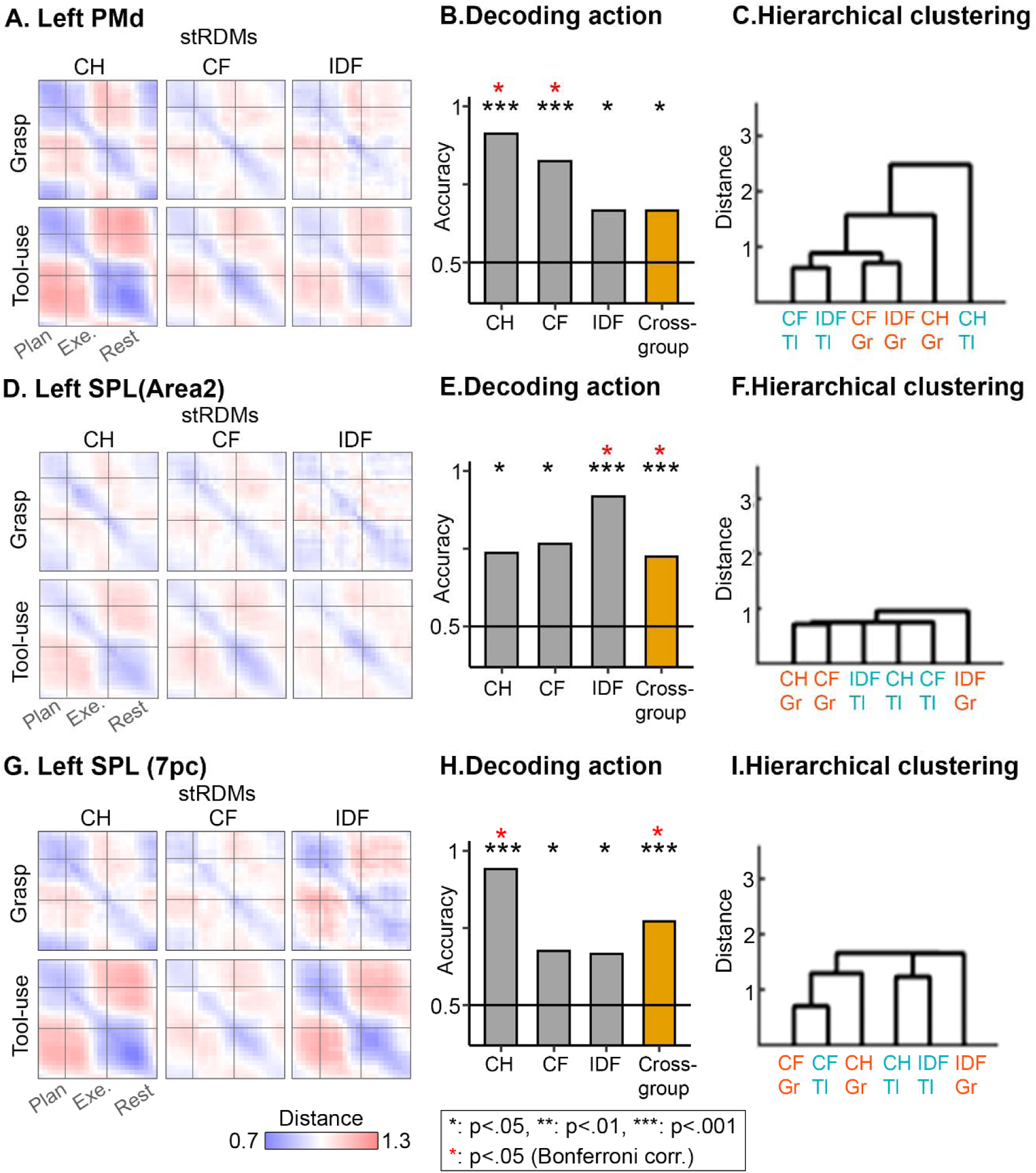
tMVPA in association tool-use areas shows action-specific features. Data is shown for the left PMd (A-C), left SPL area 2 (D-F), and left SPL 7pc (G-I). These are the same left-lateralized ROIs as in Fig. 3A and 2B. A, D, G: For each action-type and effector, a single-trial representational dissimilarity matrix (stRDM) is compute where the values are 1-Pearson’s r between the multivoxel response patterns of each pair of time points within a trial (left-most dissimilarity pattern). Different stages of the trials can be seen reflected in the dynamics, corresponding t a change between action planning and execution, as well as the end of execution and the beginning of the rest perio after each trial. B, E, H: The bar graph depicts decoding accuracy of action-type within each effector and across all three groups/effectors (CH, CF, IDF) based on stRDM. C, F, I: Hierarchical clustering depicts the grouping of stRDMs based on their similarity, with longer lines along y-axis denoting larger dissimilarity between items/clusters. All regions show a tendency to group by action-type instead of by effector.

## Discussion

An increasing amount of behavioral, electrophysiological, and neuroimaging evidence converges on the idea of a shared representation for reaching and grasping action-types that are independent of sensorimotor specifics, including effector (Gallivan et al., 2013; Liu et al., 2020; Liu et al., 2022). Yet, the extent to which (1) the brain supports an action-type representation beyond these actions, (2) the brain does so for non-ethological actions such as tool-use across somatotopically distant effectors, and (3) such representations are derived from sensorimotor experience remains poorly understood.

Here, we showed that tool-use actions are represented in motor association areas at an effector-independent level. Specifically, we found that the left PMd and SPL (areas 2 and 7pc), regions belonging to the typical tool-use network for hand actions, have a consistent preference for tool-use across controls’ hand, controls’ foot, and IDs’ foot. The multi-voxel response pattern in these regions additionally differentiates between actions across controls’ hand and foot and within IDs foot actions (in PMd and SPL area 2), lending further evidence to action-type representations that are independent of the executing effector. Finally, the temporal dynamics in motor association areas carry effector-independent action-type information. These findings argue that tool-use actions, while being newly evolved and almost exclusively associated with the hand, are effector-independent in higher-level motor association areas. In contrast, primary sensorimotor and adjacent cortices are selective for the acting body part. Temporal dynamics similarly show that the primary motor areas carry effector-specific information. Ultimately, these results make a case for an effector-independent tool-use network that is organized by action-type and primary sensorimotor regions that are organized by effector. Additionally, our findings hold for a population of individuals born without hands, suggesting that there is an action-type organization in motor association areas, and that it does not require hand sensorimotor experience to develop. Together, this supports a hierarchical organization for action representations where the acting effector is one such property that can be generalized across.

## The tool-use network contains effector-independent representations

Neuroimaging and lesion studies identified a left-lateralized frontoparietal network specialized for representing tool-use actions and tool-related knowledge. Typical regions include the inferior frontal gyrus (IFG), PMd, PMv, SMG, SPL, IPS, and middle temporal cortex (MTC; (Lewis 2016; Johnson-Frey 2004; Brandi et al., 2014). Further functional division is drawn between a more dorsal pathway connecting the SPL to the PMd that is involved in online tool-use action control and a ventral pathway including MTC, IPL, SMG, and PMV that subserves long-term conceptual knowledge about tools (Rizzolatti & Matelli, 2003; Binkofski & Buxbaum, 2013; Buxbaum et al., 2006; Buxbaum et al., 2007; Daprati & Sirigu, 2006; Johnson & Grafton, 2003; Johnson-Frey, 2004). We found that areas of the dorsal pathway, namely PMd and SPL, contain effector-independent representations, suggesting these regions may be organized by action-type. Importantly, it is unlikely these findings are driven by differences at the motor output level such as limb trajectory or movement duration because different objects (for grasping) and tool-use trajectories (i.e., turning left or right) were used to balance low-level factors and univariate and multivariate results were reproduced after accounting for movement onset and duration. Likewise, the temporal unfolding of brain responses that reflect effector-independence, characterized by the univariate time-course analysis and tMVPA, may reflect common computations underlying each action regardless of effector. For example, grasping involves calculating object size, and tool-use requires one to compute the relative position between the object and the tool (Osiurak et al., 2018; Liu et al., 2022).

Our findings extend existing ideas about the abstraction level of the tool-use network. Past studies observed a preference for pantomimed tool-use versus random gesturing in left IPS, PMd, PMv, SMG, and IFG, regardless of if the left or right hand was used (Johnson-Frey et al., 2005; Moll et al., 2000). Although this finding argues for hand-independent mechanisms, it is possible that the shared preference was driven by the semantic processing of objects involved only in the pantomiming condition and not necessarily an action representation (see also Przybylski & Króliczak, 2016; Brandi et al., 2014). Moreover, this study did not perform any direct cross-effector analyses, making it difficult to argue for generalization across the hands for tool-use actions. By using daily tools in both the grasping and tool-use conditions, we controlled the level of object semantics, thus isolating processes related to action execution. More importantly, past studies have only tested effector-independence across dominant and non-dominant hands which have homotopic cortical representations and are often coordinated in daily life. By performing MVPA across the hand and foot within controls and testing IDs, we provide further evidence for effector-independent tool-use action representations that are generalizable to a novel and inexperienced effector (the foot in controls) and are independent of hand-related motor experience (i.e., IDs; Ehrsson et al., 2003; Porro et al., 1996; Solodkin et al., 2004; Hanakawa et al., 2003; Lacourse et al., 2005; Schnitzler et al., 1997).

Yet, our data do not argue for effector-invariance in all areas of the tool-use network. Specifically, although PMv is thought to encode action goals and be involved in action execution and action understanding, we do not find it is consistently engaged in an effector-independent manner (**Figs. S2, S7, S13**). Similarly, the human tool-use region of anterior SMG (SMG area 2; Peeters et al., 2009) did not show a consistent preference for tool-use across effectors. Though this region showed a preference for tool-use over grasping in controls’ hand and IDs’ foot, there was no significant difference between the actions for controls’ foot actions (**Fig. S5**). This result suggests that SMG may encode some effector and/or experience information. One possibility is that these areas primarily encode conceptual knowledge about the object in addition to the action-type (Brandi et al., 2014; Binkofski & Buxbaum, 2013). If this is the case, our use of tools in both conditions could result in a less-differentiated response in these areas. Future studies are needed to examine the level of abstraction hierarchy in terms of action execution and semantic processing across tool-use areas to understand if the ventral tool pathway generalizes across effectors.

## Hierarchical representations in the motor system

The idea that the brain leverages a hierarchical structure to facilitate generalization in the motor domain has a long history (Lashley, 1950; Hebb, 1949; Stelmach & Diggles, 1982; Haar & Donchin, 2020; Merel et al., 2019) and has driven investigations into the organizational principle of the motor cortex (Rijntjes et al., 1999; Grafton and Hamilton, 2007) and action control (Badre & Nee, 2018). Recent studies have provided additional evidence for a hierarchical organization of reaching and grasping actions in the motor cortex. Human neuroimaging work has found shared representations across dominant and non-dominant hands for reaching and grasping in visuomotor areas such as the PMd, the parietal lobe, and preSMA (Criscimagna-Hemminger, 2003; Heed et al., 2011; Heed et al., 2016; Gallivan et al., 2013; Haar et al., 2017; Turella et al., 2020). More recently, studies have tested effector-independence for more distal effectors, the hand and foot, for pointing actions in the aIPS (Heed et al., 2016; Heed et al., 2011; Leoné et al., 2014) and for reaching and grasping actions in additional parietal areas (aIPS, pIPS, midIPS), SMA, and PMd (Liu et al., 2020). However, despite behavioral support (Osiurak et al., 2018; Rosenbaum 1980), there is a paucity of neural evidence for effector-independence for other actions, and especially for uniquely human actions.

Here, we present evidence that effector is one such property that can be generalized across for non-ethological tool-use representations. By expanding the scope of effector independence to include non-ethological actions, distal effectors, and individuals born without hands, our results further suggest that effector-independence is a general feature of action representations. Studies have shown that there are shared kinematic features and activation patterns in PMd/v and PPC between actions performed by the hand and hand-held tools (Gallivan et al., 2013; Tresilian & Stelmach, 1997; Gentilucci et al., 2004). This results in the possibility that motor association areas have co-opted effector-independent principles of ethological actions and applied them to more complex tool-use actions.

The idea that areas of the motor cortex are organized based on action-type has been shown in non-human primates, suggesting that it is a fundamental principle established through evolution. Studies applying long trains of electrical stimulation to motor regions of non-human primates have discovered an action map whereby each zone encodes for complete multi-joint sequences of ethological actions including hand-to-mouth actions, defensive movements, object manipulation actions, and reach-to-grasp actions (Graziano et al., 2002; Graziano et al., 2005; Cooke & Graziano, 2004a; 2004b; Graziano, 2006; Kaas et al., 2013). These studies provided evidence that the motor system is organized by the performed action in addition to the effector. However, long-train stimulation is limited in its ability to target specific brain areas due to propagation (Cheney et al., 2013). By leveraging the spatial accuracy of fMRI to show that action-type is the driving organizational principle in people born without hands, we extend past work by demonstrating that the action-type based organization can be effector-independent and is not limited to ethological actions.

In contrast, early areas in the motor hierarchy show selectivity for the executing body part, regardless of the action, supporting a somatotopic organization. The temporal dynamics revealed by tMVPA contain effector-specific features in primary motor areas but not association tool-use regions (**Fig. S10**). Critically, hand actions elicited greater activation in M1 hand area than for IDs despite hand function being taken over by feet in these individuals’ daily lives. These findings extend past studies on IDs when they performed simple muscle flexes (i.e. flexing/extending toes, Striem-Amit et al., 2018) and ethological actions (Liu et al., 2020), together suggesting that across motor domains, the organization principle of primary sensorimotor cortex is innately constrained by somatotopy and shows less functional plasticity compared to association cortex.

We are, however, by no means suggesting that the motor system’s organization is strictly hierarchical. Instead, the motor system may be organized compositionally such that there is a mixed code of multiple representation dimensions (Willett et al., 2020), or gradients of abstraction within an area. Our univariate analysis showed a main effect of effector and of action-type in different areas of the parietal lobe (**Figs. 1,2**) and the temporal dynamic patterns in M1 areas carry action information for non-native effectors (**Fig. 6**), suggesting information beyond one level of a hierarchical gradient. This is in accordance with evidence for co-existing representations of multiple effectors and of both effector and action-types within given motor areas (Muret et al., 2022; Graziano & Aflalo, 2007; Meier et al., 2008; Andersen and Aflalo, 2022; Diomedi et al., 2020; Nau et al., 2024; Taghizadeh et al., 2024; Tye et al., 2024; Zhang et al., 2017), proposed as an effective way of reducing dimensionality in representations (Badre and Nee, 2018; Tye et al., 2024). Studies have also proposed the idea of a modular hierarchy with separate concrete to abstract gradients within the parietal lobe, premotor cortex, and even the primary cortex (Hatsopoulos et al., 2007; Hocherman and Wise, 1991; Graziano et al., 2002a; Liu et al., 2020; Kadmon Harpaz et al., 2014; Heed et al., 2016; Leoné et al., 2014; Gordon et al., 2023) which could serve as a way to facilitate neuromotor redundancy such that the same action can be executed many different ways. The existence of a mixed code or of gradients of representations could further be a way of supplying feedback during online motor control such that even if there is a generalizable motor plan, the error corrections and feedback could still be specific to the executing body part (Kawato et al., 1987; Quirmbach & Limanowski, 2024).

While our data support a large-scale hierarchical organization in the motor domain, future studies are needed to investigate how an abstract action-type is translated into specific sensorimotor parameters through either mixed-coding at smaller scales or cortico-cortical connections.

## Plasticity principles in congenital handlessness

The fact that the tool-use network supports tool-use for the feet in people born without hands is consistent with our previous findings of ethological actions in this population. We recently showed that the dissociation between reaching and grasping, and the preference for the action-type between them develops in people born without hands (Liu et al., 2020). This suggests that the motor association areas maintain their computational roles for action regardless of the acting body part, even in people born without the primary effector (see also Hahamy et al., 2017). This is achieved through the typical generalization of this system: effector independence in the typical brain could allow for maintaining the same action even when the primary effector is missing from birth. More broadly, our findings with IDs are consistent with findings in people born blind and people born deaf. In both these cases, it appears that association cortical regions maintain their typical computations by replacing the missing input channel: visual cortex category selectivity (for script, body shapes, objects or faces) seems to develop in people born blind for the same categories perceived through audition or touch (Bi et al., 2016; Mattioni et al., 2020; Peelen et al., 2013; Pietrini et al., 2004; Striem-Amit and Amedi, 2014; Striem-Amit et al., 2012; van den Hurk et al., 2017). Similarly, the language system in congenital deafness retains its processing preferences for sign language (Corina et al., 2003; Emmorey, 2021; MacSweeney et al., 2004), and visual tasks recruit the parallel-function auditory cortex (Benetti et al., 2017; Bola et al., 2017; Lomber, 2017). In this work, we expand these findings to the tool-use network, suggesting that much like for perception, the association action system at large can maintain its functional roles regardless of sensorimotor experience. This appears to contrast with the primary motor cortex in IDs, which maintains an effector-based preference, as described above. In the context of reorganization, it reinforces the link between the original organization principles and the plasticity afforded in deprivation: association cortices whose organization is task-or domain-based but that generalize across inputs or outputs can flexibly accommodate the loss of a certain input or output channel. Primary cortices whose organization is specific to low-level properties are bound to this level of organization even in early deprivation. Together, this double dissociation in reorganization highlights the capacities of plasticity in the human brain.

## Implications for brain-machine interfaces

The concept of effector-independence has potential implications for brain-computer-interfaces (BCIs; Lebedev & Nicolelis, 2017; Gallego et al., 2022). Most current prosthetics rely on lower-level kinematic information to restore function (Lebedev & Nicolelis, 2017). However, the existence of an effector-independent abstract code and temporal dynamics makes the generalization to alternative effectors easily feasible. Indeed, learning to leverage a shared code between the hand and the foot in the premotor area facilitated motor recovery via brain-computer-interfaces in individuals with tetraplegia when compared to using effector-specific information only (Willett et al., 2020). Likewise, it is possible to record from areas of the parietal lobe to command neurprosthetics and restore motor function to those living with paralysis (Andersen et al., 2019). Similar claims have been recently made about speech rehabilitation, where neuroprosthetics may read out the desired language output even across languages (Silva et al., 2024). These support the readout of a more abstract code, and at a level of goal or action-type selection, towards driving BCIs.

## Conclusions

In conclusion, we investigated whether effector-independence extended to non-ethological tool-use actions. We found that several core regions of the tool-use network, including the PMd and areas of the SPL, had a shared preference for tool-use across the hand and the foot, that was independent of sensorimotor experience. Notably, these results were bolstered by multivariate and temporal dynamics analyses. Together, our findings point to a hierarchical representation of actions in the motor system where there is generalization across the executing effector and regardless of effector experience. This supports an organization based on action-type in the motor association cortex. Last, our findings show the link between the hierarchical organization of the typically developed cortex and the reorganization present in deprivation.

## Materials and Methods

### Participants

Twenty-three typically developed control participants (11 females; mean age: 31.05; age range: 20-65) and six individuals born with severely shortened or completely absent upper limbs (individuals with upper-limb dysplasia; abbreviated IDs; two females; mean age: 40; age range: 21-62; see **Table S5** for additional details) participated in the experiment. There was no significant difference in age between groups (U = 49.5, *p* = 0.21, non-parametric Mann-Whitney U-test). Data from several of these IDs had been reported in previous neuroimaging and behavioral studies (see **Table S6** for participant ID matching; Liu et al., 2020; Striem-Amit et al., 2018; Striem-Amit et al., 2017; Vannuscorps et al., 2019; Vannuscorps & Caramazza, 2016a; Vannuscorps & Caramazza 2016b). Importantly, all recruited IDs perform everyday actions primarily using their feet. Since most previous action-related studies use right-handed participants (Cavina-Pratesi et al., 2018; Cavina-Pratesi et al., 2010; Culham et al., 2003; Frey et al., 2005; Chapman et al., 2002; Liu et al., 2020), and the majority of ID participants are right footed, controls were required to be right-handed and right-footed (handedness/footedness was determined by self-report which has been shown to strongly correspond to experimental measures; (Coren, 1993). However, due to the rarity of complete and bilateral congenital upper-limb dysplasia, IDs were recruited regardless of foot dominance. Five IDs were right-footed, and one ID (ID6) was left-footed. This experiment focuses on actions using the dominant effector, and therefore, all participants performed the experiment with their right hands and feet, except ID6, who completed the actions with her dominant left foot. All experimental protocols and procedures were approved by the Institutional Review Board of Georgetown University in accordance with the Declaration of Helsinki. Written informed consent was collected from each participant, and they were compensated for their time.

### Experimental design and procedures

The neuroimaging experiment was a slow event-related design, with each type of effector (feet in IDs, hands and feet in controls) tested in two runs. Due to technical considerations, each run corresponded to action execution by only one effector (846TRs per run). The first four typically developed participants had an additional experimental condition which was later omitted, resulting in runs of 950TRs. Each run began with a baseline period of 7.8 seconds (10TRs), followed by pseudorandomized tool-use and grasping trials (six grasping and 12 tool-use trials per run). For each grasp action, participants reached and grasped a spatula or a crayon without lifting the object. For each tool-use action, participants reached and grasped a spatula and used it to spread balloons to the left or the right (mimicking the action of frosting a cake or spreading butter on toast; the balloons were in place also during the grasping condition). Participants began with their arms and legs resting naturally. During hand runs, participants had their legs propped up slightly so that the angle of the board on which the object was placed would be comfortable for object viewing and manipulation (**Fig. S1**). During foot runs, participants’ legs were comfortably resting on supportive pillows to facilitate viewing of foot actions and minimize upper body and head movement. Objects were placed on a “grasparatus” (Culham et al., 2003) at a comfortable distance. In both hand and foot runs an experimenter inside the scanner replaced the objects on a trial-by-trial basis. Participants were instructed to fixate on a cross located centrally on a screen viewed from a mirror mounted on the head coil between trials. Two of the IDs (ID2 and ID5) were unable to directly see the board as they lay supine, so data was collected as they viewed the objects using the mirror mounted on the head coil. These two IDs were instructed to close their eyes during action execution to avoid confusion caused by mirroring. For each grasping action, participants reached toward and grasped the object between the first and second digits of the hand or foot (thumb and index finger of the hand and big toe and second toe of the foot, respectively). Object order in a run was balanced and pseudo-randomized.

Each trial began with an auditory instruction indicating the action (e.g., “turn right”). Following the instructions, participants had 4.7 seconds (6TRs; action planning) before hearing the first sound cueing them to begin executing the action (action execution). Participants had 5.5 seconds (7TRs) to complete the action and were instructed to hold the action’s end position until they heard a second sound, cueing them to return to the resting position (return period). During the return period participants would either return the object to its starting position on the board (hand trial) or drop it (foot trial; as most control participants could not quickly return the object with their feet; 1.6 seconds, 2TRs). Trials were spaced by 8.6 seconds (11TRs). Separate practice sessions were carried out prior to the in-scanner experiment to familiarize the participants with the trial timing and object manipulation instructions.

All trials were visually inspected by an experimenter in real-time to ensure they were accurately performed. Trials that were missed or wrongly executed by the participants were excluded from further analyses. Data for four typically developed participants were excluded because of an inability to complete the tasks.

### Functional Imaging

Imaging data were collected in a Siemens Prisma-Fit 3T scanner at the Center for Functional and Molecular Imaging at Georgetown University Medical Center using a 32-channel head coil. Functional data were acquired with a T2*-weighted segmented gradient echo-planar imaging sequence (repetition time [TR]= 782 ms, echo time [TE]= 35 ms, flip angle [FA] = 52 degrees, FOV=208×200 mm, matrix = 104×100, slice thickness = 2 mm, distance factor = 0%, voxel size = 2×2×2mm^3^). The whole brain was covered by 72 interleaved slices, transversally oriented. For each participant, a T1-weighted anatomical reference volume was collected with an MPRAGE sequence (TR = 1900 ms, inversion time [TI] = 900 ms, FA = 9 degrees, FOV = 256×256mm, distance factor = 50%, 1 mm isotropic resolution). The whole brain was covered in 176 ascending slices in a sagittal orientation. Data that support the main findings of this study, as well as the source maps for the figures data have been deposited in OSF (https://osf.io/jwv9q/).

## Data Analysis

### Preprocessing

Imaging data were preprocessed and analyzed using BrainVoyager (Brain Innovation, BV22.4). Functional images were preprocessed using BrainVoyager’s standard procedures (slice scan time correction, 3D motion correction to the first volume of each run using rigid-body transformation, and high-pass temporal filtering – cutoff frequency: two cycles per scan). The first two volumes in each functional run were discarded to allow for magnetization stabilization. Functional and anatomical datasets for each participant were aligned and fit to standard Talairach space (Talairach and Tournoux, 1988). Anatomical cortical reconstruction procedures included segmentation of the white matter using a grow-region function embedded in BrainVoyager. Analyses were conducted in volumetric space and then superimposed onto cortical space.

### Additional motion correction steps

Given that hand and foot actions may induce head movements, we included additional motion correction steps:

Ventricle and White Matter Regressors – Subject-specific masks were defined separately for white matter (WM) and ventricular cerebrospinal fluid (CSF) tissues prior to any spatial smoothing using T1-weighted anatomical images. The normalized average signals of WM and CSF were included as separate regressors of no interest for each participant [Weissenbacher et al., 2009; Hallquist et al., 2013; Caballero-Gaudes & Reynolds, 2017).

Motion estimation – BrainVoyager rigid-body motion correction yields a six-dimensional time series (three translation directions and three rotation directions) which were z-normalized and used as regressors of no interest. We also included the normalized first temporal derivative of motion to explain additional variance due to motion by accounting for a one-frame delay of its effect on the BOLD signal (Satterthwaite et al., 2013; Maknojia et al., 2019; Siegel et al., 2014; Power et al., 2014).

Framewise displacement calculation and TR censoring – The framewise displacement was calculated as described in (Power et al., 2012) and individual timepoints that showed head motion exceeding 1mm were censored (Wilke & Baldeweg, 2019; Lemieux et al., 2007; Siegel et al., 2014).

Exclusion of data – Any run with more than 50 TRs censored (corresponding to approximately 5-6% of data loss) was excluded from further analysis. A total of 10 runs were excluded (four from controls’ hand data and six from controls’ foot data). No runs from IDs were excluded. One control participant’s entire dataset was excluded from analyses due to this criterion. For the remaining participants, there was no significant difference in volumes censored between groups (H(2) = 4.96, *p* = 0.08, as calculated by a Kruskal-Wallis test).

### Univariate analysis

For all univariate analyses, functional data files were spatially smoothed using a 6mm FWHM kernel. Each stage (action planning, action execution, return) of the conditions of interest (grasping, tool-use) was modeled separately. Three different general linear models (GLMs) were generated (one per group/effector) for univariate analyses, focusing on the action execution stage. Random effect (RFX) GLMs were used for the controls, separately for the hand and foot experiments; fixed-effect GLM were used for the IDs due to the sample size (Friston et al., 1999). We leveraged RFX ANOVAs to examine (1) the main effect of effector, (2) the main effect of action-type, and (3) the interaction between the two parameters. To understand the consistency across actions performed by controls with different effectors, we computed an ANOVA including controls’ hand and controls’ foot data (CH+CF, **Fig. 1C**) as a two-way ANOVA, with within-subject factors of effector (hand, foot) and action-type (grasp, tool-use). A second ANOVA was computed to inspect the effect of actions across the groups as each uses its dominant effector (controls’ hand and IDs’ foot; CH+IDFs, Fig. 1F), thus controlling for the potential confound that controls represent foot actions via manual imagery. This ANOVA included one within-subject factor (action-type) and one between-subject factor (group, implying also the use of a different effector). Activation maps were thresholded at *p* < 0.001 and corrected for multiple comparisons at *p* < 0.05 using the spatial extent method, a set-level statistical inference correction used in BrainVoyager (Forman et al., 1995; Friston et al., 1994). To examine results that correspond to a consistent action-type preference or a consistent effector preference across the two populations, the spatial overlaps between the main effects of the two ANOVAs were calculated (**Fig. 2A** for effector, **Fig. 2B** for action-type).

ANOVA main effect analyses do not inherently clarify the direction of action preference. Additionally, the inclusion of controls’ hand data in both sets of ANOVAs raises the possibility that the shared main effect could be driven by controls’ hand data. To mitigate this, results from the group-level ANOVA analyses were complemented by direct contrasts looking for tool-use preference (tool-use > grasp along with tool-use > rest, to focus on positive activation) for each group separately (**Fig. S7**). Given the stricter contrast, each activation map was thresholded at *p* < 0.01 and corrected for multiple comparisons at *p* < 0.05 as described above. The three maps were later overlayed and a spatial overlap was calculated (**Fig. 3C**). Given the use of a fixed-effects GLM in the IDs, this analysis was additionally performed in each ID to ascertain inter-individual consistency (**Fig. S8**).

To directly test whether typical tool-use regions are effector-independent, a region-of-interest (ROI) analysis was performed. ROIs were defined from the peak F-value from an ANOVA of the controls’ hand data (thresholded at *p* < 0.01 and corrected for multiple comparisons at *p* < 0.05). We focused on left cortical ROIs and right cerebellar ROIs given the typical cortical left-dominance in tool-use and action representations (Diedrichsen et al., 2009; Diedrichsen et al., 2011; Eickhoff et al., 2005; Eickhoff et al., 2006; Eickhoff et al., 2007; Johnson-Frey 2004; Kumar et al., 2020; Merrick et al., 2020) (**Tables S3 and S4**). Controls’ action-selective hand ROIs were further restricted to those with a tool-use preference. For each ROI, beta values for each group were extracted and tested for the significance of the difference between the conditions in a two-tailed paired student’s t-test (corrected for multiple comparisons using a Benjamini-Hochberg FDR correction; Benjamini and Yekutieli, 2001).

### Motor behavior recording and analysis

To verify that all main findings were not driven by a difference in either execution onset times or action duration, we replicated all core analyses while explicitly modeling the hemodynamic response with the movement onset and duration recorded with accelerometers during the scans.

During the MRI experiments, BIOPAC tri-axial TSD109C2-MRI accelerometers were used to capture movement kinematic data. An accelerometer was placed on D1 (thumb or big toe) and D2 (index finger or second toe) of the participants’ dominant hand/foot performing the actions in each run. Data were missing due to a technical error for four accelerometer runs (belonging to typically developed participants), which resulted in the exclusion of one additional subject for this control analysis. The duration-corrected analyses are based on 16 typically developed controls and six individuals born without hands.

The accelerometers recorded tri-axial data with a sampling frequency of 500Hz. Accelerometer data were analyzed using custom scripts in Python 3.11.4. The norm acceleration of the accelerometer data was calculated across both accelerometers 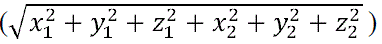 (adapted from Maimon-Mor et al., 2020; Mannini et al., 2013; Schaefer et al., 2014; van Hees et al., 2011). A 4^th^ order butterworth bandpass filter was applied to the normed accelerometer data between the frequencies of 0.2Hz and 15Hz to remove high-frequency noise and gravitational artefacts (Maimon-Mor et al., 2020; Mannini et al., 2013; Schaefer et al., 2014; van Hees et al., 2011). Velocity was calculated as the integral of the bandpassed accelerometer data. A fourth-order high-pass filter was applied at 0.05Hz to remove a low frequency drift in the velocity data. Jerk (changes in acceleration) was calculated as the derivative of the bandpassed accelerometer data. Therefore, high jerks are anticipated for abrupt changes in movement as would be expected at the beginning and the end of an action. The absolute values of velocity and jerk were used to holistically characterize movement kinematics inside the scanner.

Two independent raters made determinations on the beginning and end of movement at the single trial level. The onset of movement was determined by a sharp, high jerk. The end of the movement was based on a relative stabilization of the movement at the end of each execution stage, as participants were instructed to complete the action and then stay in the final position until the return signal. Trials were re-rated if there was a discrepancy of more than 700ms between raters’ decisions. If ratings based on two independent raters did not converge, a third rater independently arbitrated between the values. To quantitatively assess the agreement between the two raters, an intraclass correlation (ICC) was calculated using the ‘psych’ package in R (version 4.4.1). The analysis resulted in an ICC(C,1) value of 1 for both the beginning and the end of movement, indicating a near-perfect agreement between the raters (correlations between raters were 0.99 for both the beginning and end of movement). As a result, the average movement onset and end times of the two raters per trial were used to re-define the action planning and execution regressors for this control analysis. Separate linear-mixed effects models were used to assess differences in reaction time (from the auditory “go” cue to movement onset) and duration between grasping and tool-use separately per group (controls’ hand, controls’ foot, and IDs’ foot). Trial number and action-type were treated as fixed effects and subject identity was included as a random intercept to account for individual differences in baseline duration. T-tests using Satterthwaite’s method were used to calculate the significance. There were significant differences between the durations of the actions across all three groups (controls’ hand: t(491.403) = 20.41, *p* < 0.001; controls’ foot: t(465.271) = 18.160, *p* < 0.001; IDs’ foot: t(198.007) = 15.75, *p* < 0.001), with tool-use taking longer. Only controls’ hand had a significant difference between the reaction times of action (controls’ hand: t(490.766) = –2.32, *p* = 0.021; controls’ foot: t(465.159) = –1.896, *p* = 0.059; IDs’ foot: t(197.993) = 0.05, *p* = 0.96) (see **Fig. S11** for graphs showing the mean durations and reaction times).

### Multivariate analyses

For all multivariate analyses, functional data were spatially smoothed using a 3mm FWHM kernel.

*ROI MVPA.* ROI MVPA was performed for tool-use selective ROIs defined using univariate results (areas identified in **Fig. 3A,B**) using a linear support vector machine classifier implemented in the CoSMoMVPA toolbox (Oosterhof, et al., 2016). GLM beta estimates were obtained for each trial using SPM12. Given additional head movement and potential noise in the data, which could artificially increase the decoding accuracy, we baselined our responses to a control region in addition to baselining to chance level (50%). The white matter within the bilateral anterior temporal lobe (ATL) was chosen because we do not expect it to contain information about the execution of the actions. Given that the paradigm involved an auditory instruction and visuo-motor task, few areas could serve as control regions. Ventricle regions were not used as controls because of the implicit masking implemented in SPM. We defined two 4mm radius spheres to match the relative size of the tool-use ROIs. We performed subject-specific baselining of the decoding in each ROI by subtracting from it the decoding accuracy from the bilateral ATL.

Given the imbalance in trial number between the grasping and the tool-use trials, conditions are balanced prior to decoding, resulting in several tool-use trials being discarded. To achieve a robust and fair estimate of the accuracy values, each baselined decoding accuracy was computed 1,000 times, discarding different trials at random. During each iteration, the ROI decoding accuracy was calculated along with the ATL decoding accuracy using the same training and testing partitions. To report only results with greater decoding in the ROI than in both ATL and chance level, the maximum between the ATL decoding accuracy and chance (max(0.5, ATL_accuracy)) was taken as the baselining value for that iteration, and this was subtracted from the ROI decoding accuracy. The average of a subject’s 1,000 baselined iterations was stored. To evaluate significance, a one-tailed, paired t-test against zero was performed at the for each group and effector of interest. Significance is reported at *p* < 0.05, and FDR-corrected across ROIs for each effector or group.

### Cross-effector whole-brain MVPA

To test for a shared action representation between the hand and foot beyond these tool-use areas, we additionally performed a whole-brain searchlight MVPA within the gray matter between grasping and tool-use across controls’ hand and feet. The analysis was performed using a linear support vector machine classifier implemented in the CoSMoMVPA toolbox (Oosterhof, et al., 2016) with each spherical searchlight containing 200 voxels. Classification accuracies were calculated using cross-validation such that the classifier was trained on hand runs and then tested on foot runs and vice-versa. The results from the two folds were averaged per participant. Searchlight decoding was computed 100 times per participant to ensure stability of accuracy values. One-sample *t*-tests against chance level (50%) were performed at group level across the brain. To evaluate significance, 100 null maps were generated for each subject by shuffling action-type labels across trials. These null maps were then randomly sampled for 10,000 iterations to perform multiple-comparison correction at the group level using the Monte Carlo cluster statistics in CoSMoMVPA (Oosterhof, et al., 2016). Group-level results were thresholded at *p* < 0.0001 and corrected for multiple comparisons at *p* < 0.05.

### Temporal analyses methods

*Time-course GLM analysis.* To test if the temporal dynamics of the action network are similar for hand and foot actions, we leveraged the identical trial order across body parts (with the exception of control participants C1-C4, where an additional condition was included). We used the time courses from the controls’ hand data as predictors in a GLM of the IDs foot data. Average time courses across the controls hand runs were sampled from one effector-independent region (PMd; defined from the CH+CF action-type effect ANOVA) and one effector-dependent region (hand M1; defined from the CH+CF effector effect ANOVA) for this analysis. GLM calculations and contrasts were computed as described above.

### Temporal multivariate pattern analysis (tMVPA)

Temporal multivariate pattern analysis was performed in accordance with Vizioli et al. (2018). The preprocessed time courses were divided by the temporal mean and segmented into trials of 26TRs from the beginning of the auditory cue to the last TR of the inter-trial baseline. For each participant and ROI, a single-trial representational dissimilarity matrix (stRDM) was calculated for each combination of effector and action-type. Specifically, the 1 – Pearson’s correlation was calculated between the response pattern at each time point in Trial X with each time point in Trial Y, forming a 26-by-26 RDM between Trial X and Trial Y. This process was repeated for all possible pairs of trials, and the resulting RDMs were averaged as a final stRDM for each individual. Finally, the stRDMs were averaged across participants, resulting in one stRDM for each effector/group and action-type (e.g. **Fig 6A**). As with Vizioli et al., we performed multivariate analyses on the Fisher z-transformed correlation coefficients and plotted the (1 – raw correlation coefficient) for visualization. ROIs used were those defined in the univariate ROI analysis (tool-use effector-independent regions) and hand and foot M1 ROIs defined from the CH+CF effector effect ANOVA.

Multivariate pattern analysis was performed on the stRDMs to test whether each effector and action is associated with distinct temporal dynamics patterns (**Fig. 6 middle panels**). The within-effector decoding of action-type was performed in a leave-one-subject-out manner. In each iteration, an SVM classifier was trained on the grasping and tool-use stRDMs from all but one participant and tested on the left-out participant. The accuracies from all iterations were then averaged. For the decoding of effector/group within action, in each iteration an stRDM from controls’ hand, controls’ foot, and IDs’ foot was randomly left out and a three-way SVM classifier was trained on the remaining data. The accuracies from all iterations were then averaged. The binary decoding analyses were implemented in CoSMoMVPA, and the three-way decoding analyses were implemented using the *fitcecoc* function in MATLAB.

We also decoded action across effectors/groups and decoded effectors/groups across actions. For cross-effector/group decoding of action-type, an SVM classifier was trained on the grasping and tool-use stRDMs of participants from one effector/group in each iteration (e.g., controls’ hand), and tested on the other two effector/groups (e.g., controls’ foot and IDs’ foot). For cross-action decoding of effector/group, in each iteration a three-way SVM classifier was trained on stRDMs from one action-type to differentiate controls’ hand, controls’ foot, and IDs’ foot, and tested on the other action-type.

We computed permutation tests to determine the significance of the decoding performance at group level. For the decoding of action-type, we generated 100 pairs of null grasping and tool-use stRDMs for each effector in each participant by shuffling the action-type label across trials (i.e., 100 pairs of random maps for each control’s hand, each control’s foot, and each IDs’ foot). In each permutation iteration, a pair of null stRDMs were randomly drawn from each participant, forming a “null group” based on which a decoding accuracy was obtained. The accuracies over 1000 permutation iterations form a null distribution, among which the percentage of accuracies greater than the actual decoding accuracy was the permutation p-value. For the decoding of effector within and across action-type, permutation tests were performed by shuffling the effector labels across participants 1000 times. Permutation p-values were calculated as the percentage of null accuracies greater than the actual decoding accuracy.

To rule out the possibility that differences in stRDM are driven by head motion signals, we constructed stRDMs based on the six head motion parameters. For example, the six parameters at time point 1 in Trial X were correlated with the six parameters at time point 2 in Trial Y and converted to dissimilarity. We then performed multivariate pattern analyses as described above on the head motion stRDMs to examine if each action-type is associated with a unique head motion temporal pattern.

Finally, hierarchical clustering analyses were performed based on the Euclidean distance between stRDMs using the *linkage* function in MATLAB.

## Acknowledgements

We are thankful to the participants who participated in our experiment. We thank Nanak Nihal Khalsa for his contribution to the collection of the data. This work was supported by the Edwin H. Richard and Elisabeth Richard von Matsch Distinguished Professorship in Neurological Diseases (to E.S.A.) and a Shanghai Youth Science and Technology Innovation Plan (22YF1454200 to Y.L.).

## Notes

### Competing Interest Statement

The authors have declared no competing interest.

## References

1. Alexander, G. E., & Crutcher, M. D. (1990). Neural representations of the target (goal) of visually guided arm movements in three motor areas of the monkey. Journal of neurophysiology, 64(1), 164–178.

2. Amunts, K., Mohlberg, H., Bludau, S., & Zilles, K. (2020). Julich-Brain: A 3D probabilistic atlas of the human brain’s cytoarchitecture. Science, 369(6506), 988–992.

3. Andersen, R.A., and Aflalo, T. (2022). Preserved cortical somatotopic and motor representations in tetraplegic humans. Current Opinion in Neurobiology 74, 102547.

4. Andersen, R. A., Aflalo, T., & Kellis, S. (2019). From thought to action: The brain–machine interface in posterior parietal cortex. Proceedings of the National Academy of Sciences, 116(52), 26274–26279.

5. Badre, D., and Nee, D.E. (2018). Frontal Cortex and the Hierarchical Control of Behavior. Trends in Cognitive Sciences 22, 170–188.

6. Benjamini, Y., & Yekutieli, D. (2001). The control of the false discovery rate in multiple testing under dependency. Annals of statistics, 1165–1188.

7. Benetti, S., van Ackeren, M.J., Rabini, G., Zonca, J., Foa, V., Baruffaldi, F., Rezk, M., Pavani, F., Rossion, B., and Collignon, O. (2017). Functional selectivity for face processing in the temporal voice area of early deaf individuals. Proceedings of the National Academy of Sciences.

8. Bi, Y., Wang, X., and Caramazza, A. (2016). Object Domain and Modality in the Ventral Visual Pathway. Trends in Cognitive Sciences.

9. Bola, Ł., Zimmermann, M., Mostowski, P., Jednoróg, K., Marchewka, A., Rutkowski, P., and Szwed, M. (2017). Task-specific reorganization of the auditory cortex in deaf humans. Proceedings of the National Academy of Sciences.

10. Brandi, M. L., Wohlschläger, A., Sorg, C., & Hermsdörfer, J. (2014). The neural correlates of planning and executing actual tool use. Journal of Neuroscience, 34(39), 13183–13194.

11. Binkofski, F., & Buxbaum, L. J. (2013). Two action systems in the human brain. Brain and language, 127(2), 222–229.

12. Bonferroni, C. (1936). Teoria statistica delle classi e calcolo delle probabilita. Pubblicazioni del R istituto superiore di scienze economiche e commericiali di firenze, 8, 3–62.

13. Buxbaum, L. J., Kyle, K. M., Tang, K., & Detre, J. A. (2006). Neural substrates of knowledge of hand postures for object grasping and functional object use: Evidence from fMRI. Brain research, 1117(1), 175–185.

14. Buxbaum, L. J., Kyle, K., Grossman, M., & Coslett, B. (2007). Left inferior parietal representations for skilled hand-object interactions: evidence from stroke and corticobasal degeneration. Cortex, 43(3), 411–423.

15. Caballero-Gaudes, C., & Reynolds, R. C. (2017). Methods for cleaning the BOLD fMRI signal. Neuroimage, 154, 128–149.

16. Cavina-Pratesi, C., Connolly, J. D., Monaco, S., Figley, T. D., Milner, A. D., Schenk, T., & Culham, J. C. (2018). Human neuroimaging reveals the subcomponents of grasping, reaching and pointing actions. Cortex, 98, 128–148.

17. Cavina-Pratesi, C., Monaco, S., Fattori, P., Galletti, C., McAdam, T. D., Quinlan, D. J., Goodale, M., & Culham, J. C. (2010). Functional magnetic resonance imaging reveals the neural substrates of arm transport and grip formation in reach-to-grasp actions in humans. Journal of Neuroscience, 30(31), 10306–10323.

18. Chapman, H., Gavrilescu, M., Wang, H., Kean, M., Egan, G., & Castiello, U. (2002). Posterior parietal cortex control of reach to grasp movements in humans. European Journal of Neuroscience, 15(12), 2037–2042.

19. Cheney, P.D., Griffin, D.M., and Van Acker, G.M., 3rd (2013). Neural hijacking: action of high-frequency electrical stimulation on cortical circuits. Neuroscientist 19, 434–441.

20. Cisek, P., Crammond, D. J., & Kalaska, J. F. (2003). Neural activity in primary motor and dorsal premotor cortex in reaching tasks with the contralateral versus ipsilateral arm. Journal of neurophysiology, 89(2), 922–942.

21. Coren, S. (1993). Measurement of handedness via self-report: the relationship between brief and extended inventories. Perceptual and motor skills, 76(3), 1035–1042.

22. Corina, D.P., San Jose-Robertson, L., Guillemin, A., High, J., and Braun, A.R. (2003). Language lateralization in a bimanual language. J Cogn Neurosci 15, 718–730.

23. Cooke, D. F., & Graziano, M. S. (2004). Sensorimotor integration in the precentral gyrus: polysensory neurons and defensive movements. Journal of neurophysiology, 91(4), 1648–1660.

24. Criscimagna-Hemminger, S. E., Donchin, O., Gazzaniga, M. S., & Shadmehr, R. (2003). Learned dynamics of reaching movements generalize from dominant to nondominant arm. Journal of neurophysiology, 89(1), 168–176.

25. Culham, J. C., Danckert, S. L., Souza, J. F. D., Gati, J. S., Menon, R. S., & Goodale, M. A. (2003). Visually guided grasping produces fMRI activation in dorsal but not ventral stream brain areas. Experimental brain research, 153, 180–189.

26. Daprati, E., & Sirigu, A. (2006). How we interact with objects: learning from brain lesions. Trends in cognitive sciences, 10(6), 265–270.

27. Diedrichsen, J., Balsters, J. H., Flavell, J., Cussans, E., & Ramnani, N. (2009). A probabilistic MR atlas of the human cerebellum. neuroimage, 46(1), 39–46.

28. Diedrichsen, J., Maderwald, S., Küper, M., Thürling, M., Rabe, K., Gizewski, E. R., … & Timmann, D. (2011). Imaging the deep cerebellar nuclei: a probabilistic atlas and normalization procedure. Neuroimage, 54(3), 1786–1794.

29. Diomedi, S., Vaccari, F.E., Filippini, M., Fattori, P., and Galletti, C. (2020). Mixed Selectivity in Macaque Medial Parietal Cortex during Eye-Hand Reaching. iScience 23, 101616.

30. Emmorey, K. (2021). New Perspectives on the Neurobiology of Sign Languages. Frontiers in Communication 6.

31. Ehrsson, H. H., Geyer, S., & Naito, E. (2003). Imagery of voluntary movement of fingers, toes, and tongue activates corresponding body-part-specific motor representations. Journal of neurophysiology.

32. Eickhoff, S. B., Stephan, K. E., Mohlberg, H., Grefkes, C., Fink, G. R., Amunts, K., & Zilles, K. (2005). A new SPM toolbox for combining probabilistic cytoarchitectonic maps and functional imaging data. Neuroimage, 25(4), 1325–1335.

33. Eickhoff, S. B., Heim, S., Zilles, K., & Amunts, K. (2006). Testing anatomically specified hypotheses in functional imaging using cytoarchitectonic maps. Neuroimage, 32(2), 570–582.

34. Eickhoff, S. B., Paus, T., Caspers, S., Grosbras, M. H., Evans, A. C., Zilles, K., & Amunts, K. (2007). Assignment of functional activations to probabilistic cytoarchitectonic areas revisited. Neuroimage, 36(3), 511–521.

35. Emmorey, K. (2021). New perspectives on the neurobiology of sign languages. Frontiers in communication, 6, 748430.

36. Evarts, E. V. (1968). Relation of pyramidal tract activity to force exerted during voluntary movement. Journal of neurophysiology, 31(1), 14–27.

37. Forman, S. D., Cohen, J. D., Fitzgerald, M., Eddy, W. F., Mintun, M. A., & Noll, D. C. (1995). Improved assessment of significant activation in functional magnetic resonance imaging (fMRI): Use of a cluster size threshold. Magnetic Resonance in medicine, 33(5), 636–647.

38. Frey, S. H., Vinton, D., Norlund, R., & Grafton, S. T. (2005). Cortical topography of human anterior intraparietal cortex active during visually guided grasping. Cognitive Brain Research, 23(2-3), 397–405.

39. Friston, K. (2008). Hierarchical models in the brain. PLoS computational biology, 4(11), e1000211.

40. Friston, K. J., Worsley, K. J., Frackowiak, R. S., Mazziotta, J. C., & Evans, A. C. (1994). Assessing the significance of focal activations using their spatial extent. Human brain mapping, 1(3), 210–220.

41. Friston, K.J., Holmes, A.P., and Worsley, K.J. (1999). How many subjects constitute a study? Neuroimage 10, 1–5.

42. Gallego, J. A., Perich, M. G., Miller, L. E., & Solla, S. A. (2017). Neural manifolds for the control of movement. Neuron, 94(5), 978–984.

43. Gallego, J. A., Makin, T. R., & McDougle, S. D. (2022). Going beyond primary motor cortex to improve brain–computer interfaces. Trends in neurosciences, 45(3), 176–183.

44. Gallivan, J. P., McLean, D. A., Flanagan, J. R., & Culham, J. C. (2013). Where one hand meets the other: limb-specific and action-dependent movement plans decoded from preparatory signals in single human frontoparietal brain areas. Journal of Neuroscience, 33(5), 1991–2008.

45. Gallivan, Jason P., et al. “Decoding effector-dependent and effector-independent movement intentions from human parieto-frontal brain activity.” Journal of Neuroscience 31.47 (2011): 17149–17168.

46. Gallivan, Jason P., Ingrid S. Johnsrude, and J. Randall Flanagan. “Planning ahead: object-directed sequential actions decoded from human frontoparietal and occipitotemporal networks.” Cerebral Cortex 26.2 (2016): 708–730.

47. Gallivan, Jason P., et al. “Decoding the neural mechanisms of human tool use.” elife 2 (2013): e00425.

48. Giszter, S. F. (2015). Motor primitives—new data and future questions. Current opinion in neurobiology, 33, 156–165.

49. Goodall, J. (1964). Tool-using and aimed throwing in a community of free-living chimpanzees. Nature, 201(4926), 1264–1266.

50. Gordon, E. M., Chauvin, R. J., Van, A. N., Rajesh, A., Nielsen, A., Newbold, D. J., … & Dosenbach, N. U. (2023). A somato-cognitive action network alternates with effector regions in motor cortex. Nature, 617(7960), 351–359.

51. Grafton, S. T., & Hamilton, A. F. D. C. (2007). Evidence for a distributed hierarchy of action representation in the brain. Human movement science, 26(4), 590–616.

52. Gentilucci, M., Roy, A. C., & Stefanini, S. (2004). Grasping an object naturally or with a tool: are these tasks guided by a common motor representation?. Experimental Brain Research, 157(4), 496–506.

53. Graziano, M. S. (2016). Ethological action maps: a paradigm shift for the motor cortex. Trends in cognitive sciences, 20(2), 121–132.

54. Graziano, M. S., Taylor, C. S., & Moore, T. (2002). Complex movements evoked by microstimulation of precentral cortex. Neuron, 34(5), 841–851.

55. Graziano, M. S., Aflalo, T. N., & Cooke, D. F. (2005). Arm movements evoked by electrical stimulation in the motor cortex of monkeys. Journal of neurophysiology, 94(6), 4209–4223.

56. Cooke, D. F., & Graziano, M. S. (2004). Super-flinchers and nerves of steel: defensive movements altered by chemical manipulation of a cortical motor area. Neuron, 43(4), 585–593.

57. Graziano, M. (2006). The organization of behavioral repertoire in motor cortex. Annu. Rev. Neurosci., 29(1), 105–134.

58. Graziano, M. S., & Aflalo, T. N. (2007). Rethinking cortical organization: moving away from discrete areas arranged in hierarchies. The Neuroscientist, 13(2), 138–147.

59. Haar, S., Donchin, O., & Dinstein, I. (2017). Individual movement variability magnitudes are explained by cortical neural variability. Journal of Neuroscience, 37(37), 9076–9085.

60. Haar, S., & Donchin, O. (2020). A revised computational neuroanatomy for motor control. Journal of Cognitive Neuroscience, 32(10), 1823–1836.

61. Hahamy, A., Macdonald, S.N., van den Heiligenberg, F., Kieliba, P., Emir, U., Malach, R., Johansen-Berg, H., Brugger, P., Culham, J.C., and Makin, T.R. (2017). Representation of multiple body parts in the missing hand territory of congenital onehanders. Current Biology 27, 1350–1355.

62. Hallquist, M. N., Hwang, K., & Luna, B. (2013). The nuisance of nuisance regression: spectral misspecification in a common approach to resting-state fMRI preprocessing reintroduces noise and obscures functional connectivity. Neuroimage, 82, 208–225.

63. Hanakawa, T., Immisch, I., Toma, K., Dimyan, M. A., Van Gelderen, P., & Hallett, M. (2003). Functional properties of brain areas associated with motor execution and imagery. Journal of neurophysiology, 89(2), 989–1002.

64. Hatsopoulos, N. G., Xu, Q., & Amit, Y. (2007). Encoding of movement fragments in the motor cortex. Journal of Neuroscience, 27(19), 5105–5114.

65. Hebb, D. O. (1949). The first stage of perception: growth of the assembly. The Organization of Behavior, 4(60), 78–60.

66. Heed, T., Beurze, S. M., Toni, I., Röder, B., & Medendorp, W. P. (2011). Functional rather than effector-specific organization of human posterior parietal cortex. Journal of Neuroscience, 31(8), 3066–3076.

67. Heed, T., Leone, F. T., Toni, I., & Medendorp, W. P. (2016). Functional versus effector-specific organization of the human posterior parietal cortex: revisited. Journal of Neurophysiology, 116(4), 1885–1899.

68. Hocherman, S., & Wise, S. P. (1991). Effects of hand movement path on motor cortical activity in awake, behaving rhesus monkeys. Experimental brain research, 83, 285–302.

69. Holdefer, R. N., & Miller, L. E. (2002). Primary motor cortical neurons encode functional muscle synergies. Experimental Brain Research, 146, 233–243.

70. Ishibashi, R., Pobric, G., Saito, S., & Lambon Ralph, M. A. (2016). The neural network for tool-related cognition: an activation likelihood estimation meta-analysis of 70 neuroimaging contrasts. Cognitive Neuropsychology, 33(3-4), 241–256.

71. Johnson-Frey, S. H. (2004). The neural bases of complex tool use in humans. Trends in cognitive sciences, 8(2), 71–78.

72. Johnson, S. H., & Grafton, S. T. (2003). From ‘acting on’to ‘acting with’: the functional anatomy of object-oriented action schemata. Progress in brain research, 142, 127–139.

73. Johnson-Frey, S. H., Newman-Norlund, R., & Grafton, S. T. (2005). A distributed left hemisphere network active during planning of everyday tool use skills. Cerebral cortex, 15(6), 681–695.

74. Kaas, J. H., Gharbawie, O. A., & Stepniewska, I. (2013). Cortical networks for ethologically relevant behaviors in primates. American journal of primatology, 75(5), 407–414.

75. Kadmon Harpaz, N., Flash, T., & Dinstein, I. (2014). Scale-invariant movement encoding in the human motor system. Neuron, 81(2), 452–462.

76. Kakei, S., Hoffman, D. S., & Strick, P. L. (2001). Direction of action is represented in the ventral premotor cortex. Nature neuroscience, 4(10), 1020–1025.

77. Kakei, S., Hoffman, D. S., & Strick, P. L. (2003). Sensorimotor transformations in cortical motor areas. Neuroscience research, 46(1), 1–10.

78. Kawato, M., Furukawa, K., & Suzuki, R. (1987). A hierarchical neural-network model for control and learning of voluntary movement. Biological cybernetics, 57, 169–185.

79. Krubitzer, L. A., & Prescott, T. J. (2018). The combinatorial creature: cortical phenotypes within and across lifetimes. Trends in neurosciences, 41(10), 744–762.

80. Kumar, A., Panthi, G., Divakar, R., & Mutha, P. K. (2020). Mechanistic determinants of effector-independent motor memory encoding. Proceedings of the National Academy of Sciences, 117(29), 17338–17347.

81. Lacourse, M. G., Orr, E. L., Cramer, S. C., & Cohen, M. J. (2005). Brain activation during execution and motor imagery of novel and skilled sequential hand movements. Neuroimage, 27(3), 505–519.

82. Lashley, K. S. (1950). In search of the engram.

83. Lebedev, M. A., & Nicolelis, M. A. (2017). Brain-machine interfaces: from basic science to neuroprostheses and neurorehabilitation. Physiological reviews, 97(2), 767–837.

84. Lemieux, L., Salek-Haddadi, A., Lund, T. E., Laufs, H., & Carmichael, D. (2007). Modelling large motion events in fMRI studies of patients with epilepsy. Magnetic resonance imaging, 25(6), 894–901.

85. Leo, A., Handjaras, G., Bianchi, M., Marino, H., Gabiccini, M., Guidi, A., Scilingo, E.P., Pietrini, P., Bicchi, A., Santello, M., & Ricciardi, E. (2016). A synergy-based hand control is encoded in human motor cortical areas. Elife, 5, e13420.

86. Leoné, F. T., Heed, T., Toni, I., & Medendorp, W. P. (2014). Understanding effector selectivity in human posterior parietal cortex by combining information patterns and activation measures. Journal of Neuroscience, 34(21), 7102–7112.

87. Lewis, J. W. (2006). Cortical networks related to human use of tools. The neuroscientist, 12(3), 211–231.

88. Liu, Y., Vannuscorps, G., Caramazza, A., & Striem-Amit, E. (2020). Evidence for an effector-independent action system from people born without hands. Proceedings of the National Academy of Sciences, 117(45), 28433–28441.

89. Liu, Y., Caracoglia, J., Sen, S., Freud, E., & Striem-Amit, E. (2022). Are reaching and grasping effector-independent? Similarities and differences in reaching and grasping kinematics between the hand and foot. Experimental Brain Research, 240(6), 1833–1848.

90. Lomber, S.G. (2017). What is the function of auditory cortex when it develops in the absence of acoustic input? Cognitive Development.

91. MacSweeney, M., Campbell, R., Woll, B., Giampietro, V., David, A.S., McGuire, P.K., Calvert, G.A., and Brammer, M.J. (2004). Dissociating linguistic and nonlinguistic gestural communication in the brain. Neuroimage 22, 1605–1618.

92. Magri, C., Fabbri, S., Caramazza, A., & Lingnau, A. (2019). Directional tuning for eye and arm movements in overlapping regions in human posterior parietal cortex. Neuroimage, 191, 234–242.

93. Maimon-Mor, R. O., Obasi, E., Lu, J., Odeh, N., Kirker, S., MacSweeney, M., … & Makin, T. R. (2020). Talking with your (artificial) hands: Communicative hand gestures as an implicit measure of embodiment. IScience, 23(11).

94. Maknojia, S., Churchill, N. W., Schweizer, T. A., & Graham, S. J. (2019). Resting state fMRI: Going through the motions. Frontiers in neuroscience, 13, 825.

95. Mannini, A., Intille, S. S., Rosenberger, M., Sabatini, A. M., & Haskell, W. (2013). Activity recognition using a single accelerometer placed at the wrist or ankle. Medicine and science in sports and exercise, 45(11), 2193.

96. Mattioni, S., Rezk, M., Battal, C., Bottini, R., Cuculiza Mendoza, K.E., Oosterhof, N.N., and Collignon, O. (2020). Categorical representation from sound and sight in the ventral occipito-temporal cortex of sighted and blind. eLife 9, e50732.

97. Meier, J. D., Aflalo, T. N., Kastner, S., & Graziano, M. S. (2008). Complex organization of human primary motor cortex: a high-resolution fMRI study. Journal of neurophysiology, 100(4), 1800–1812.

98. Merel, J., Botvinick, M., & Wayne, G. (2019). Hierarchical motor control in mammals and machines. Nature communications, 10(1), 1–12.

99. Merrick, C. M., Dixon, T. C., Breska, A., Lin, J., Chang, E. F., King-Stephens, D., Laxer, K.D., Weber, P.B., Carmena, J., Knight, R.T., & Ivry, R. B. (2022). Left hemisphere dominance for bilateral kinematic encoding in the human brain. elife, 11, e69977.

100. Moll, J., de Oliveira-Souza, R., Passman, L. J., Cunha, F. C., Souza-Lima, F., & Andreiuolo, P. A. (2000). Functional MRI correlates of real and imagined tool-use pantomimes. Neurology, 54(6), 1331–1336.

101. Muret, D., Root, V., Kieliba, P., Clode, D., & Makin, T. R. (2022). Beyond body maps: Information content of specific body parts is distributed across the somatosensory homunculus. Cell reports, 38(11).

102. Mushiake, H., Inase, M., & Tanji, J. (1990). Selective coding of motor sequence in the supplementary motor area of the monkey cerebral cortex. Experimental brain research, 82, 208–210.

103. Nau, M., Schmid, A.C., Kaplan, S.M., Baker, C.I., and Kravitz, D.J. (2024). Centering cognitive neuroscience on task demands and generalization. Nature Neuroscience.

104. Oosterhof, N. N., Connolly, A. C., & Haxby, J. V. (2016). CoSMoMVPA: multi-modal multivariate pattern analysis of neuroimaging data in Matlab/GNU Octave. Frontiers in neuroinformatics, 10, 27.

105. Osiurak, F., Jarry, C., & Le Gall, D. (2010). Grasping the affordances, understanding the reasoning: toward a dialectical theory of human tool use. Psychological review, 117(2), 517.

106. Osiurak, F., Navarro, J., Reynaud, E., & Thomas, G. (2018). Tools don’t—and won’t—make the man: A cognitive look at the future. Journal of Experimental Psychology: General, 147(5), 782.

107. Orban, G. A., & Caruana, F. (2014). The neural basis of human tool use. Frontiers in psychology, 5, 310.

108. Overduin, S. A., d’Avella, A., Roh, J., Carmena, J. M., & Bizzi, E. (2015). Representation of muscle synergies in the primate brain. Journal of Neuroscience, 35(37), 12615–12624.

109. Peelen, M.V., Bracci, S., Lu, X., He, C., Caramazza, A., and Bi, Y. (2013). Tool Selectivity in Left Occipitotemporal Cortex Develops without Vision. Journal of Cognitive Neuroscience, 1–10.

110. Peeters, R., Simone, L., Nelissen, K., Fabbri-Destro, M., Vanduffel, W., Rizzolatti, G., & Orban, G. A. (2009). The representation of tool use in humans and monkeys: common and uniquely human features. Journal of Neuroscience, 29(37), 11523–11539.

111. Penfield, W., & Boldrey, E. (1937). Somatic motor and sensory representation in the cerebral cortex of man as studied by electrical stimulation. Brain, 60(4), 389–443.

112. Pietrini, P., Furey, M.L., Ricciardi, E., Gobbini, M.I., Wu, W.H., Cohen, L., Guazzelli, M., and Haxby, J.V. (2004). Beyond sensory images: Object-based representation in the human ventral pathway. Proc Natl Acad Sci U S A 101, 5658–5663. Epub 2004 Apr 5652.

113. Poggio, T., & Bizzi, E. (2004). Generalization in vision and motor control. Nature, 431(7010), 768–774.

114. Povinelli, D. (2003). Folk physics for apes: The chimpanzee’s theory of how the world works. Oxford University Press.

115. Porro, C. A., Francescato, M. P., Cettolo, V., Diamond, M. E., Baraldi, P., Zuiani, C., Massimo, B., & Di Prampero, P. E. (1996). Primary motor and sensory cortex activation during motor performance and motor imagery: a functional magnetic resonance imaging study. Journal of Neuroscience, 16(23), 7688–7698.

116. Power, J. D., Mitra, A., Laumann, T. O., Snyder, A. Z., Schlaggar, B. L., & Petersen, S. E. (2014). Methods to detect, characterize, and remove motion artifact in resting state fMRI. Neuroimage, 84, 320–341.

117. Power, J. D., Barnes, K. A., Snyder, A. Z., Schlaggar, B. L., & Petersen, S. E. (2012). Spurious but systematic correlations in functional connectivity MRI networks arise from subject motion. Neuroimage, 59(3), 2142–2154.

118. Przybylski, Ł., & Króliczak, G. (2017). Planning functional grasps of simple tools invokes the hand-independent praxis representation network: an fMRI study. Journal of the International Neuropsychological Society, 23(2), 108–120.

119. Quirmbach, F., & Limanowski, J. (2024). Visuomotor prediction during action planning in the human frontoparietal cortex and cerebellum. Cerebral Cortex, 34(9), bhae382.

120. Reynaud, E., Lesourd, M., Navarro, J., & Osiurak, F. (2016). On the neurocognitive origins of human tool use: A critical review of neuroimaging data. Neuroscience & Biobehavioral Reviews, 64, 421–437.

121. Rijntjes, M., Dettmers, C., Büchel, C., Kiebel, S., Frackowiak, R. S., & Weiller, C. (1999). A blueprint for movement: functional and anatomical representations in the human motor system. Journal of Neuroscience, 19(18), 8043–8048.

122. Rizzolatti, G., Luppino, G., & Matelli, M. (1998). The organization of the cortical motor system: new concepts. Electroencephalography and clinical neurophysiology, 106(4), 283–296.

123. Rizzolatti, G., & Matelli, M. (2003). Two different streams form the dorsal visual system: anatomy and functions. Experimental brain research, 153, 146–157.

124. Rosenbaum, D. A. (1980). Human movement initiation: specification of arm, direction, and extent. Journal of Experimental Psychology: General, 109(4), 444.

125. Satterthwaite, T. D., Elliott, M. A., Gerraty, R. T., Ruparel, K., Loughead, J., Calkins, M. E., Eickhoff, S.B., Hakonarson, H., Gur, R.C., Gur, R.E., & Wolf, D. H. (2013). An improved framework for confound regression and filtering for control of motion artifact in the preprocessing of resting-state functional connectivity data. Neuroimage, 64, 240–256.

126. Schaefer, C. A., Nigg, C. R., Hill, J. O., Brink, L. A., & Browning, R. C. (2014). Establishing and evaluating wrist cutpoints for the GENEActiv accelerometer in youth. Medicine and science in sports and exercise, 46(4), 826.

127. Schnitzler, A., Salenius, S., Salmelin, R., Jousmäki, V., & Hari, R. (1997). Involvement of primary motor cortex in motor imagery: a neuromagnetic study. Neuroimage, 6(3), 201–208.

128. Seed, A., & Byrne, R. (2010). Animal tool-use. Current biology, 20(23), R1032–R1039.

129. Siegel, J. S., Power, J. D., Dubis, J. W., Vogel, A. C., Church, J. A., Schlaggar, B. L., & Petersen, S. E. (2014). Statistical improvements in functional magnetic resonance imaging analyses produced by censoring high motion data points. Human brain mapping, 35(5), 1981–1996.

130. Silva, A. B., Liu, J. R., Metzger, S. L., Bhaya-Grossman, I., Dougherty, M. E., Seaton, M. P., … & Chang, E. F. (2024). A bilingual speech neuroprosthesis driven by cortical articulatory representations shared between languages. Nature Biomedical Engineering, 1–15.

131. Solodkin, A., Hlustik, P., Chen, E. E., & Small, S. L. (2004). Fine modulation in network activation during motor execution and motor imagery. Cerebral cortex, 14(11), 1246–1255.

132. Stelmach, G. E., & Diggles, V. A. (1982). Control theories in motor behavior. Acta psychologica, 50(1), 83–105.

133. Striem-Amit, E., Vannuscorps, G., & Caramazza, A. (2017). Sensorimotor-independent development of hands and tools selectivity in the visual cortex. Proceedings of the National Academy of Sciences, 114(18), 4787–4792.

134. Striem-Amit, E., Vannuscorps, G., & Caramazza, A. (2018). Plasticity based on compensatory effector use in the association but not primary sensorimotor cortex of people born without hands. Proceedings of the National Academy of Sciences, 115(30), 7801–7806.

135. Striem-Amit, E., and Amedi, A. (2014). Visual cortex extrastriate body-selective area activation in congenitally blind people “seeing” by using sounds. Curr Biol 24, 687–692.

136. Striem-Amit, E., Cohen, L., Dehaene, S., and Amedi, A. (2012). Reading with sounds: sensory substitution selectively activates the visual word form area in the blind. Neuron 76, 640–652.

137. Taghizadeh, B., Fortmann, O., and Gail, A. (2024). Position– and scale-invariant object-centered spatial localization in monkey frontoparietal cortex dynamically adapts to cognitive demand. Nature Communications 15, 3357.

138. Talairach, J. & Torunoux, P. (1988). Co-planar stereotaxic atlas of the human brain.

139. Turella, L., Rumiati, R., & Lingnau, A. (2020). Hierarchical action encoding within the human brain. Cerebral cortex, 30(5), 2924–2938.

140. Tye, K.M., Miller, E.K., Taschbach, F.H., Benna, M.K., Rigotti, M., and Fusi, S. (2024). Mixed selectivity: Cellular computations for complexity. Neuron.

141. van den Hurk, J., Van Baelen, M., and Op de Beeck, H.P. (2017). Development of visual category selectivity in ventral visual cortex does not require visual experience. Proceedings of the National Academy of Sciences 114, E4501–E4510.

142. van Hees, V. T., Renström, F., Wright, A., Gradmark, A., Catt, M., Chen, K. Y., … & Franks, P. W. (2011). Estimation of daily energy expenditure in pregnant and non-pregnant women using a wrist-worn tri-axial accelerometer. PloS one, 6(7), e22922.

143. Vannuscorps, G., Andres, M., & Pillon, A. (2014). Is motor knowledge part and parcel of the concepts of manipulable artifacts? Clues from a case of upper limb aplasia. Brain and Cognition, 84(1), 132–140.

144. Vannuscorps, G., F Wurm, M., Striem-Amit, E., & Caramazza, A. (2019). Large-scale organization of the hand action observation network in individuals born without hands. Cerebral Cortex, 29(8), 3434–3444.

145. Vannuscorps, G., & Caramazza, A. (2019). Conceptual processing of action verbs with and without motor representations. Cognitive neuropsychology, 36(7-8), 301–312.

146. Vannuscorps, G., & Caramazza, A. (2016). Typical action perception and interpretation without motor simulation. Proceedings of the National Academy of Sciences, 113(1), 86–91.

147. Vannuscorps, G., & Caramazza, A. (2016). The origin of the biomechanical bias in apparent body movement perception. Neuropsychologia, 89, 281–286.

148. Vizioli, L., Bratch, A., Lao, J., Ugurbil, K., Muckli, L., & Yacoub, E. (2018). Temporal multivariate pattern analysis (tMVPA): A single trial approach exploring the temporal dynamics of the BOLD signal. Journal of neuroscience methods, 308, 74–87.

149. Wilke, M., & Baldeweg, T. (2019). A multidimensional artefact-reduction approach to increase robustness of first-level fMRI analyses: Censoring vs. interpolating. Journal of Neuroscience Methods, 318, 56–68.

150. Windischberger, C., Langenberger, H., Sycha, T., Tschernko, E. M., Fuchsjäger-Mayerl, G., Schmetterer, L., & Moser, E. (2002). On the origin of respiratory artifacts in BOLD-EPI of the human brain. Magnetic resonance imaging, 20(8), 575–582.

151. Zhang, C.Y., Aflalo, T., Revechkis, B., Rosario, E.R., Ouellette, D., Pouratian, N., and Andersen, R.A. (2017). Partially Mixed Selectivity in Human Posterior Parietal Association Cortex. Neuron 95, 697–708.e694.

