## Supplementary Information for "Tool-use brain representations are independent of the acting body part and motor experience"

### Supplementary Figures

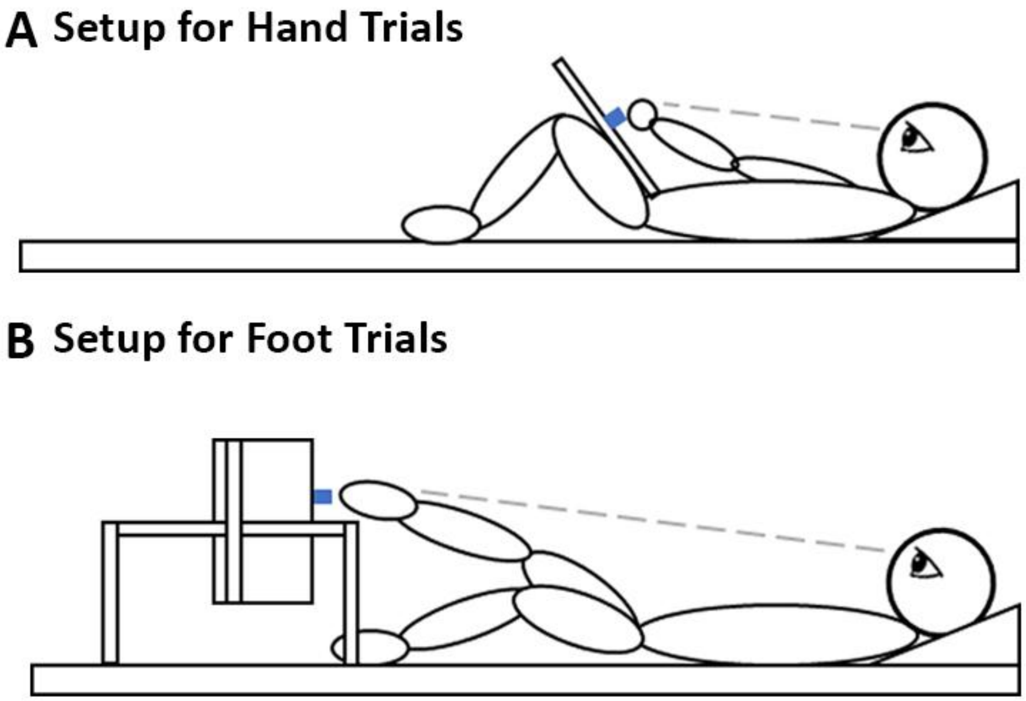


**Figure. S1. Schematic of Participant Setup and Configuration.** Participants inside the MRI scanner were auditorily cued to either grasp a spatula or a pen or turn a spatula to the right/left to mimic frosting a cake (simple tool-use). (A) Setup for participant hand trials. Participants had a board placed on their lap with an object centrally located. There was a foam pad in between their lap and the board to avoid somatosensation when changing out an object. Their knees were bent so that they could easily reach the object. (B) Setup for participant foot trials. Participants had their legs in a relaxed position with the grasparatus holding the object at a comfortable distance.

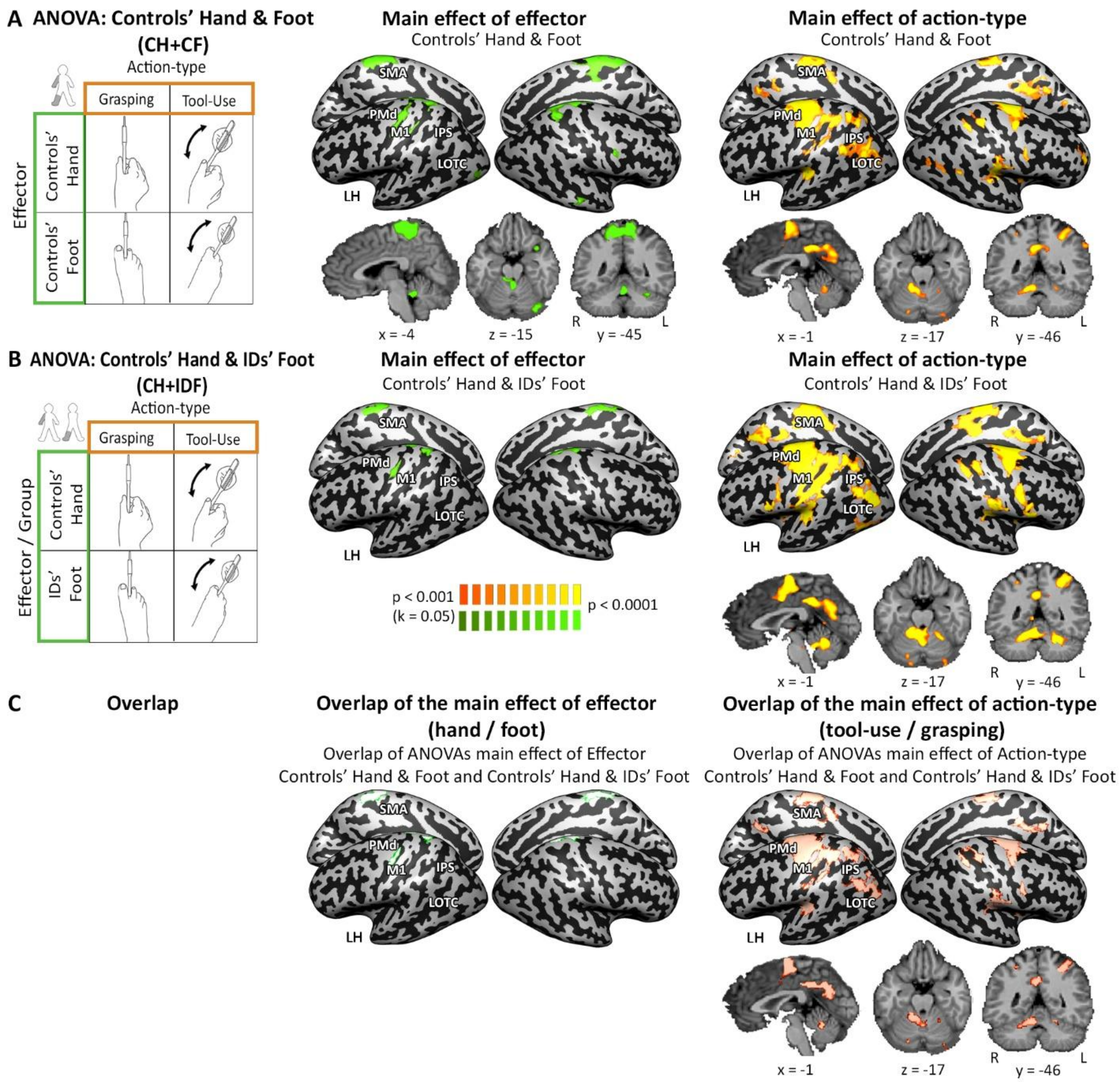


**Figure. S2. Replication of Figs. 1,2 Using Duration-Controlled Data Reveals a Consistent Frontoparietal Network involved in Action Preferences.** To control for differences in action duration between simple tool-use and grasping, we used independent motor behavior kinematic measurements for accelerometers. Results from RFX ANOVA analyses for controls' hand and controls' foot as well as controls' hand and IDs foot. Each map was thresholded at $p < 0.001$ and corrected for multiple comparisons at $p < 0.05$. (A and B left panel) Two RFX ANOVAs were performed. (A, left panel) One ANOVA was within-group and consisted of controls' hand and controls' foot data. In this ANOVA, action-type and effector were within-subjects factors. (B, left panel) The second ANOVA was across groups and consisted of controls' hand IDs foot data. In this ANOVA, effector was a between-subjects factor and action-type was a within-subjects factor. (A and B middle panel) Main effect of effector. (A, middle panel) Difference between hand and foot actions in controls was found in left M1 (foot and hand area) and SPL. (B, middle panel) Difference between hand actions in controls and foot actions in IDs (right) was found left M1 (foot area) and the SPL. (C, middle panel) Overlap of main effects of action-type shown in (A and B, middle panels). (A and B, right panel) Main effect of action-type. For controls' hand and foot (A, right panel), a shared preference for action-type independent of executing effector was found in bilateral PMd, IPS, SMA, and LOTC as well as areas left V and left VI of the cerebellum. For controls' hand and IDs foot (B, right panel) a shared preference for action-type independent of executing effector and sensorimotor experience was found in bilateral PMd, IPS, SMA, and LOTC as well as areas left V, VI, and VIIIb of the cerebellum. (C, right panel) Overlap of main effects of action-type shown in (A and B, right panels).

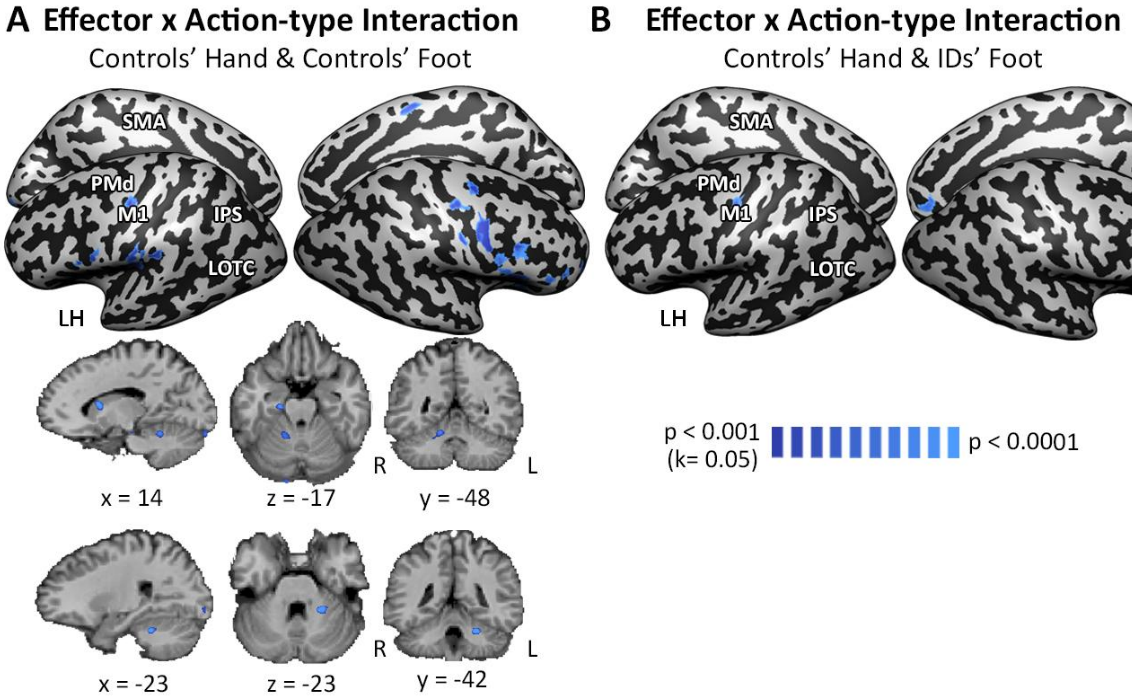


**Figure. S3. Interaction Effect between Effector and Action-type.** Using two separate RFX ANOVAs between Controls Hand and Controls Foot (CH+CF) and Controls' Hand and IDs' Foot (CH+IDF), the interaction between body part and action-type was calculated. Each map was thresholded at $p < 0.001$ and corrected for multiple comparisons at $p < 0.05$. (A) Results from CH+CF ANOVA. In This ANOVA, action-type and effector were within-subjects factors. In the left hemisphere, an area within M1 and a frontal area on the right hemisphere have an interaction effect along with two clusters in the cerebellum (right lobule IV and left lobule V). (B) Results from the CH+IDF ANOVA. This ANOVA was performed across groups and has effector as a between-subjects factor and action-type as a within-subjects factor. There is an interaction effect in an area within M1 in the left hemisphere. Notably, the area in M1 between both ANOVAs is shared and is inferior to hand M1.

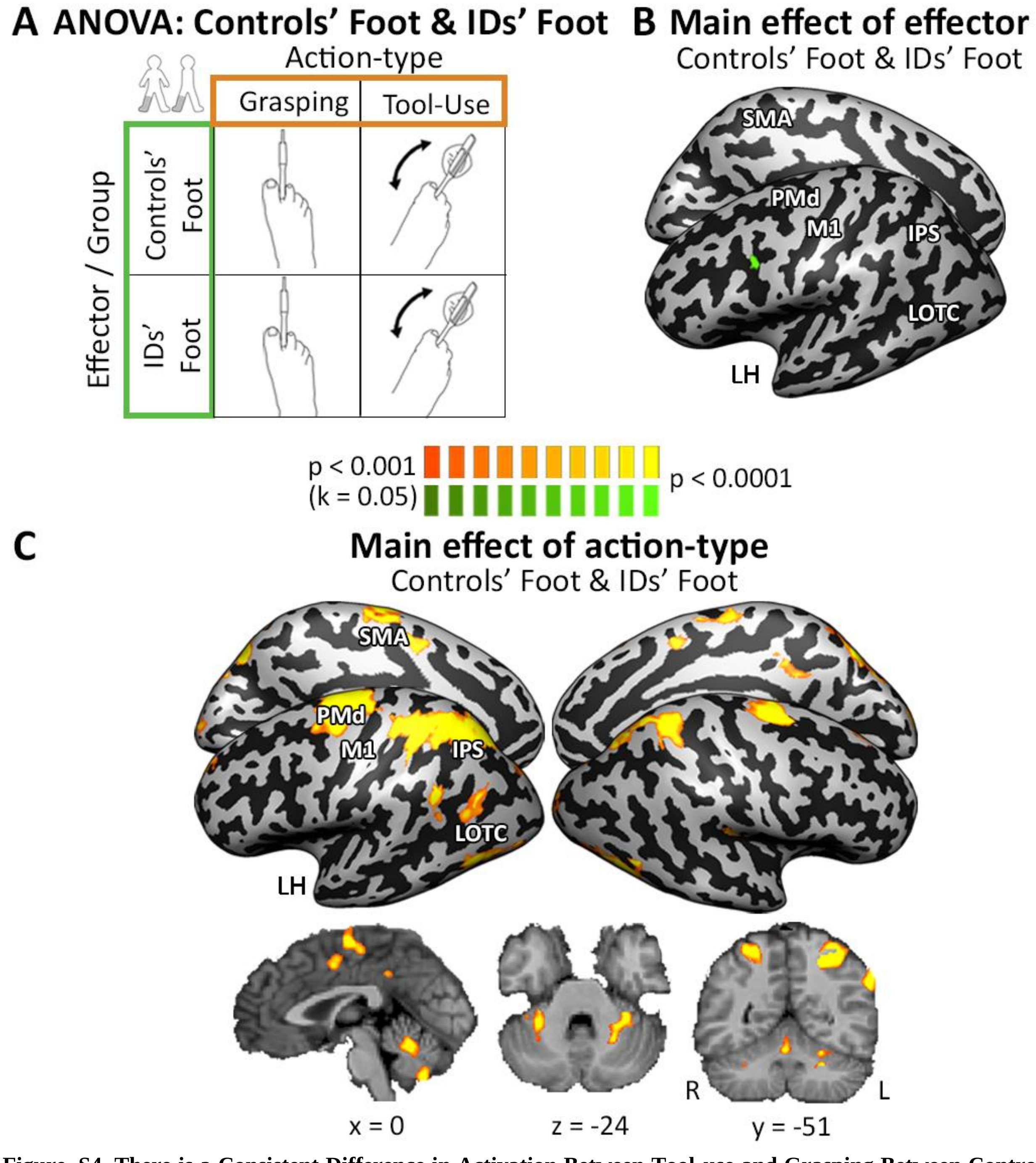


**Figure. S4. There is a Consistent Difference in Activation Between Tool-use and Grasping Between Controls Foot and IDs'.** Results from RFX ANOVA analyses between controls' foot and IDs' foot. Each map was thresholded at $p < 0.001$ and corrected for multiple comparisons at $p < 0.05$. (A) RFX ANOVA performed using effector (compensatory or atypical for IDs and controls' foot, respectively) as a between-subjects factor. (B) Difference between the foot as a compensatory or atypical effector seen in a left premotor area. (C) Shared preference for action-type independent of compensatory or atypical effector was found in the bilateral PMd, IPS, and LOTC as well as left lobule I-IV, left lobule IX, and bilateral lobule VI.

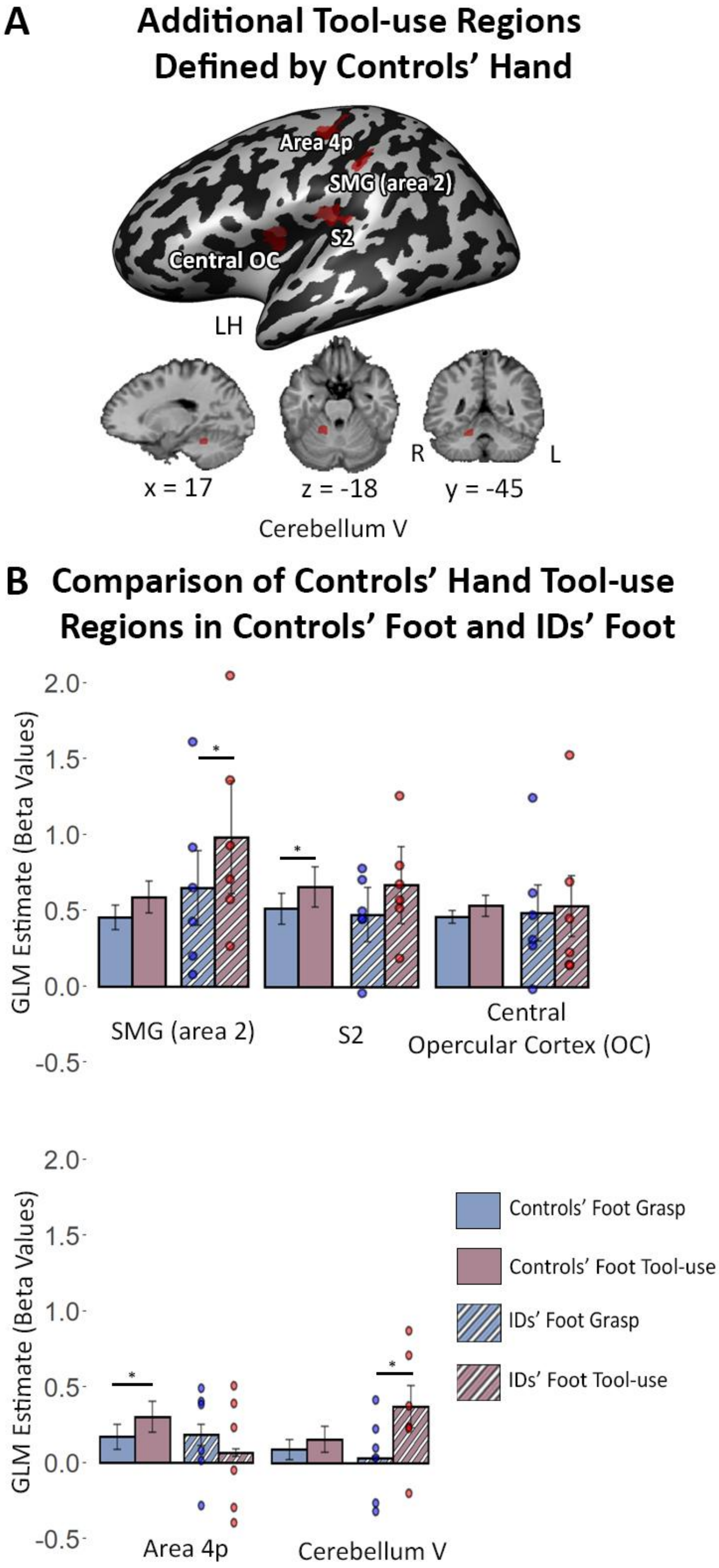
A
Additional Tool-use Regions Defined by Controls' Hand
Area 4p
SMG (area 2)
S2
Central OC
LH
R
L
x = 17
z = -18
y = -45
Cerebellum V
B
Comparison of Controls' Hand Tool-use Regions in Controls' Foot and IDs' Foot
GLM Estimate (Beta Values)
2.0
1.5
1.0
0.5
0.0
-0.5
SMG (area 2)
S2
Central Opercular Cortex (OC)
Area 4p
Cerebellum V
Controls' Foot Grasp
Controls' Foot Tool-use
IDs' Foot Grasp
IDs' Foot Tool-use

**Figure. S5. Additional Regions with Preference for Tool-use as Defined by Controls' Hand Data.** These are additional regions defined as hand tool-use areas (in addition to Fig. 2A). They do not have a consistent preference for tool-use for foot actions from the IDs and the controls' foot data (map was thresholded at $p < 0.01$ and corrected for multiple comparisons). Bar graphs of average beta values for the tool-use ROIs shown in the whole-brain for controls' foot (solid bars) and IDs (striped bars). Each dot above a striped bar plot represents one individual with dysplasia. *: $p < 0.05$. Information about these ROIs is presented in Table S2. Regions that show a preference for grasping over tool-use for controls' hand data are shown in Fig. S16. SMG: Supramarginal gyrus. Central OC: Central Opercular Cortex

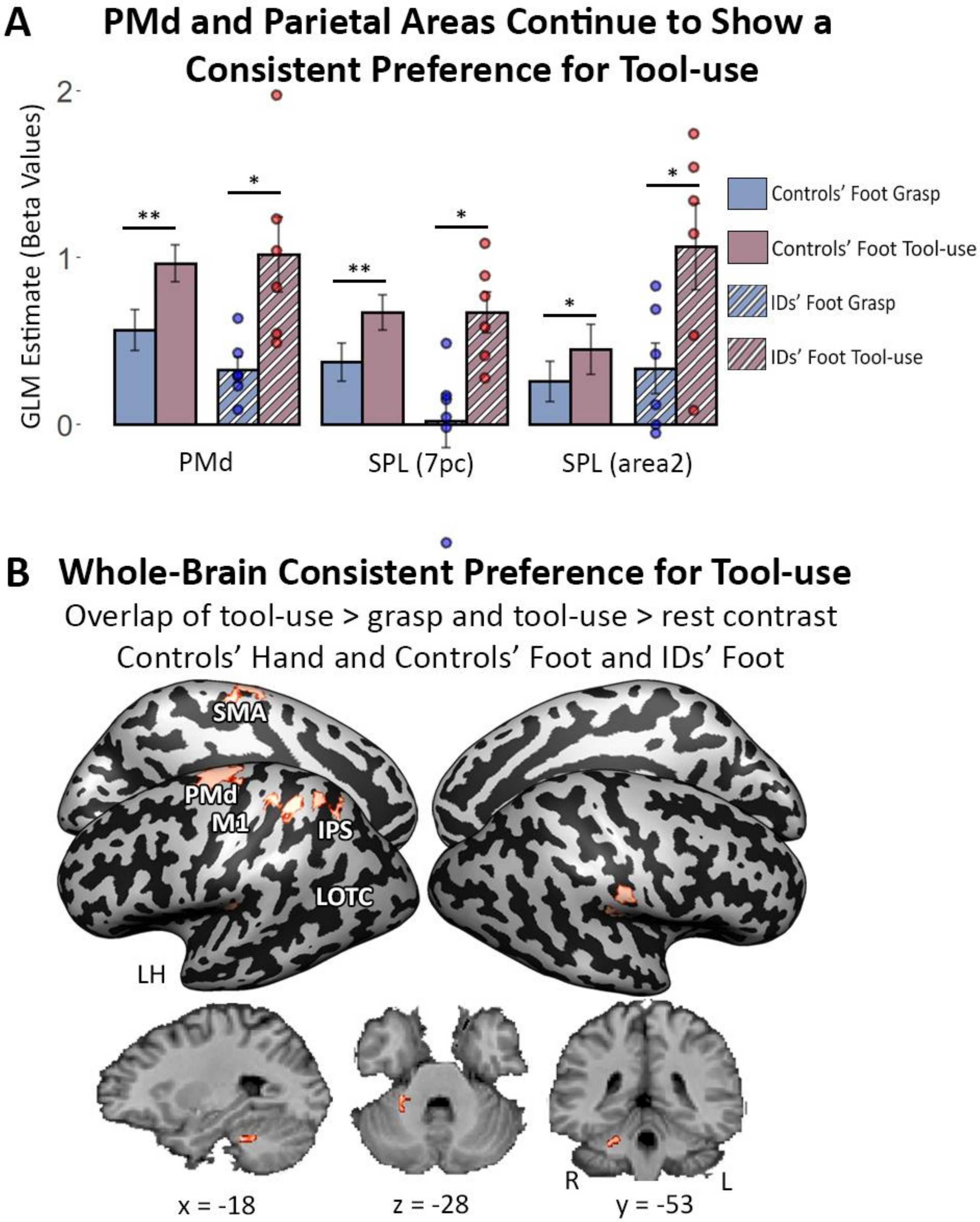


**Figure. S6. Replication of Fig. 2 Reveals Consistent Preference for Tool-use Regions Even with Duration-Controlled Data.** (A) To verify the consistent preference for tool-use over grasping seen in regions identified in Fig. 2A,B was not due to differences in action duration, we replicated the analysis in these same regions using the duration-controlled data. We show that the PMd and parietal areas continue to have a consistent preference for tool-use over grasping. Bar graphs of average beta values for the tool-use ROIs shown in (Fig. 2A) for controls' foot (solid bars)

and IDs (striped bars). Each dot above a striped bar plot represents one individual with dysplasia. The blue color corresponds to grasp and the red color corresponds to tool-use. * = $p < 0.05$; ** = $p < 0.01$, FDR-corrected. (B) We similarly repeated the whole-brain consistent tool-use preference across controls' hand, controls' foot and IDs' foot. We further show that the tool-use preference in SMA, IPS, and PMd is not due to differences in action duration (see Fig. S17 for individual group and individual ID replication). As before, each individual map was thresholded at p < 0.01 and then corrected for multiple comparisons at $p < 0.05$.

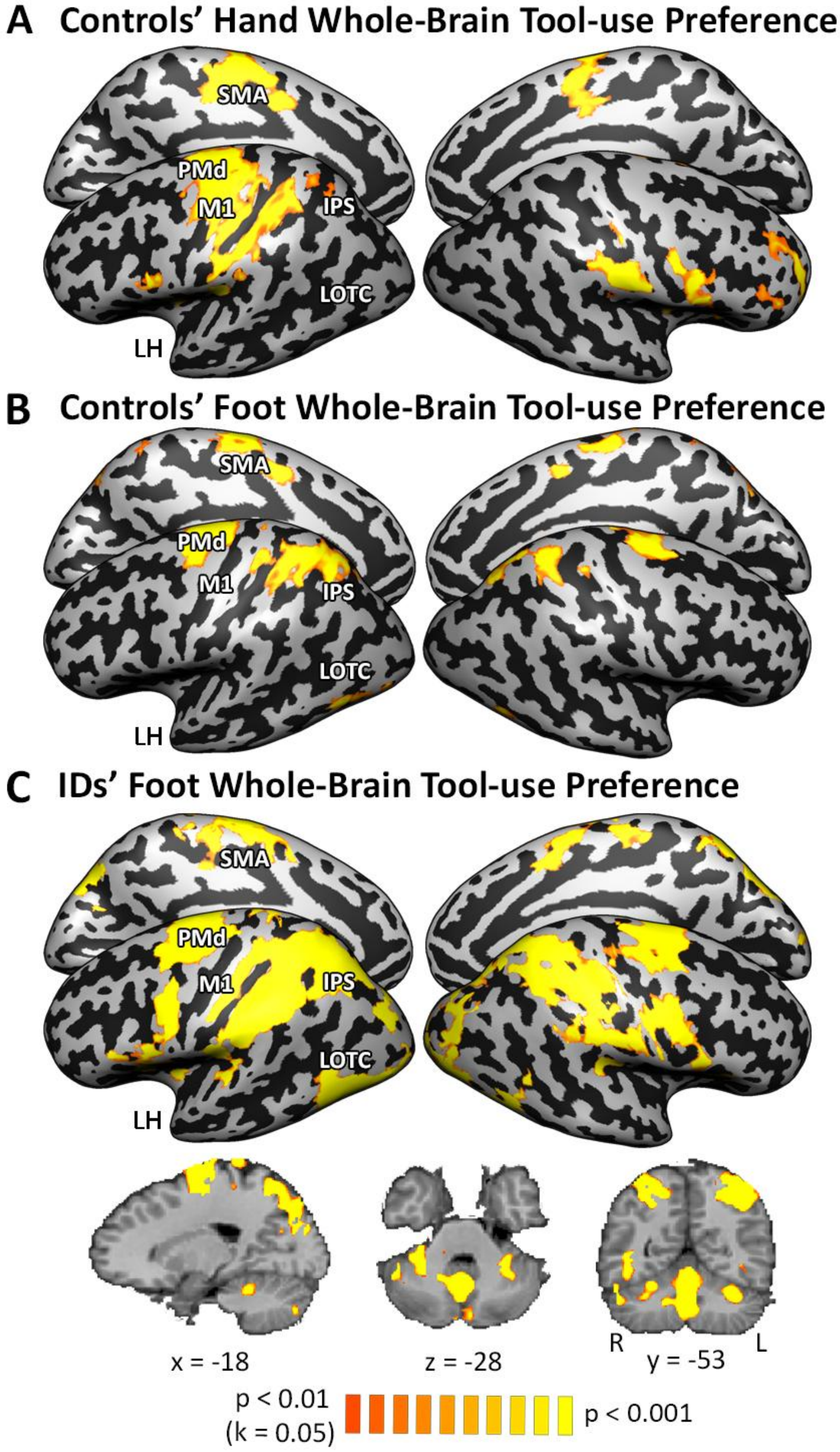

A Controls' Hand Whole-Brain Tool-use Preference
SMA
PMd
M1
IPS
LOTC
LH
B Controls' Foot Whole-Brain Tool-use Preference
SMA
PMd
M1
IPS
LOTC
LH
C IDs' Foot Whole-Brain Tool-use Preference
SMA
PMd
M1
IPS
LOTC
LH
x = -18
z = -28
R
L
y = -53
p < 0.01
(k = 0.05)
p < 0.001

**Figure. S7. Consistent, Whole-brain Tool-use Preference Across the Groups**. Individual maps of (A) controls' hand, (B) controls' foot, and (C) IDs' foot group data that were used in the whole-brain overlap contrast seen in Fig. 2C. Each map was thresholded at $p < 0.001$ and corrected for multiple comparisons at $p < 0.05$.

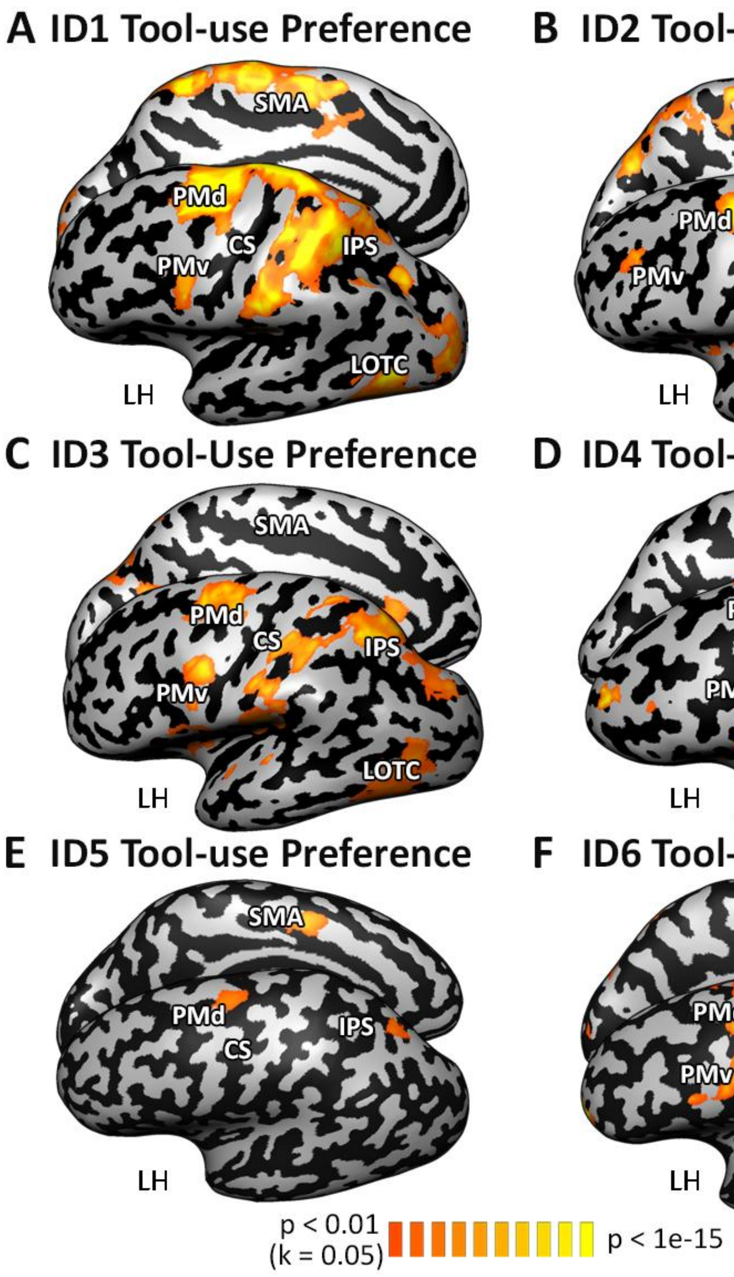


**Figure. S8. Consistent, Tool-use Preference in PMd Across All IDs**. To verify that no one ID was driving the whole-brain tool-use pattern seen in Fig. S7, we examined each IDs' contrast on their own individual brain. All IDs show a preference for tool-use over grasping in the PMd.

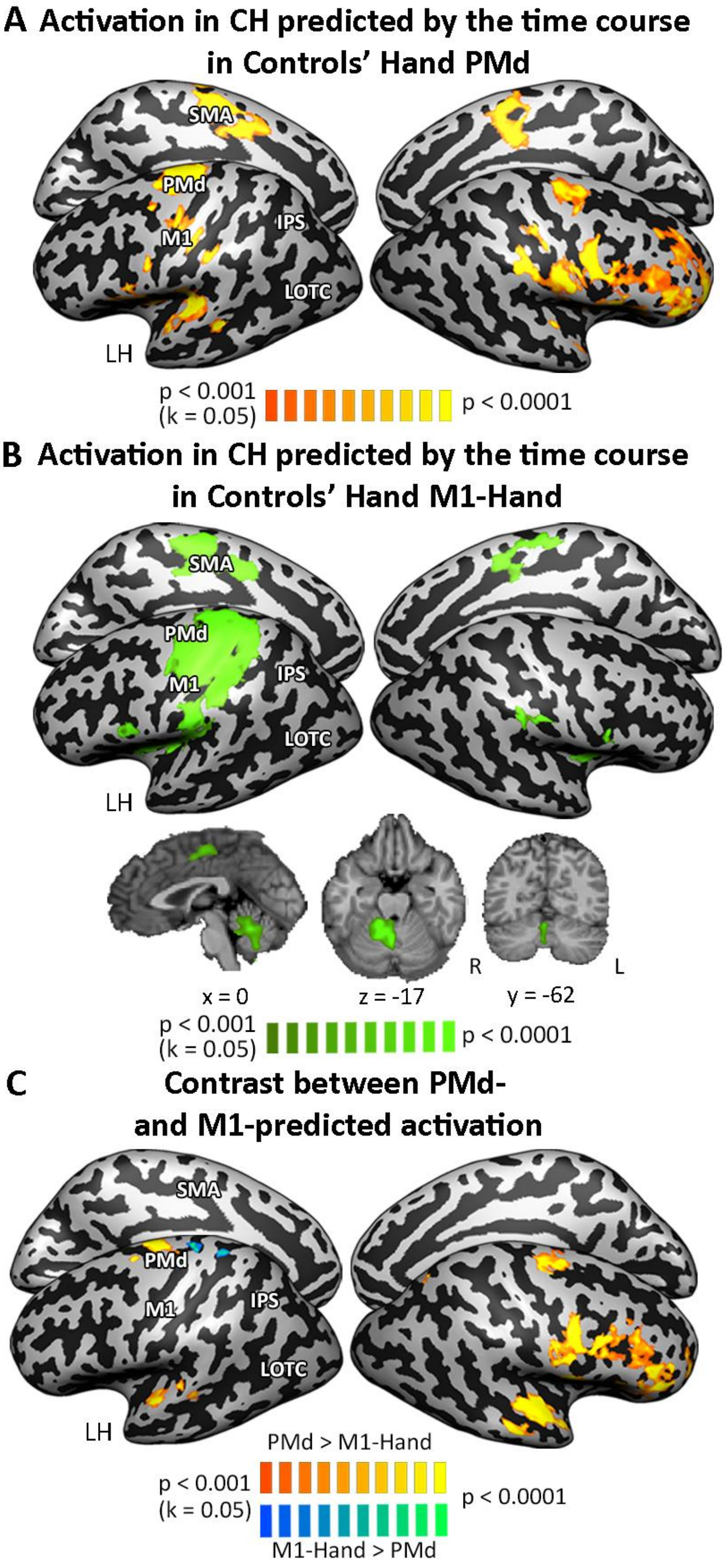
A Activation in CH predicted by the time course in Controls' Hand PMd
SMA
PMd
M1
IPS
LOTC
LH
p < 0.001 (k = 0.05)
p < 0.0001
B Activation in CH predicted by the time course in Controls' Hand M1-Hand
SMA
PMd
M1
IPS
LOTC
LH
R
L
x = 0
z = -17
y = -62
p < 0.001 (k = 0.05)
p < 0.0001
C Contrast between PMd- and M1-predicted activation
SMA
PMd
M1
IPS
LOTC
LH
PMd > M1-Hand
p < 0.001 (k = 0.05)
p < 0.0001
M1-Hand > PMd

**Figure. S9. Time-Courses from Controls' Hand Effector-Independent and Effector-Dependent Regions Reveal Effector-Independent and Effector-Dependent Networks**. All maps were thresholded at $p < 0.001$ and corrected for multiple comparisons at $p < 0.05$. (A) Using PMd (as used in Fig. 5A) as a seed region, we used its time course from controls' hand data to identify other effector-independent regions. The white line outlines regions defined as effector-independent based on the shared main effect of action-type seen in Fig. 2B. (B) Using hand M1 (as used in Fig. 5B) as a seed region, we used its time course from controls' hand data to identify other effector-dependent regions. (C) A contrast of effector-independent and effector-dependent networks highlights regions that are activated concurrently and uniquely with each of the networks.

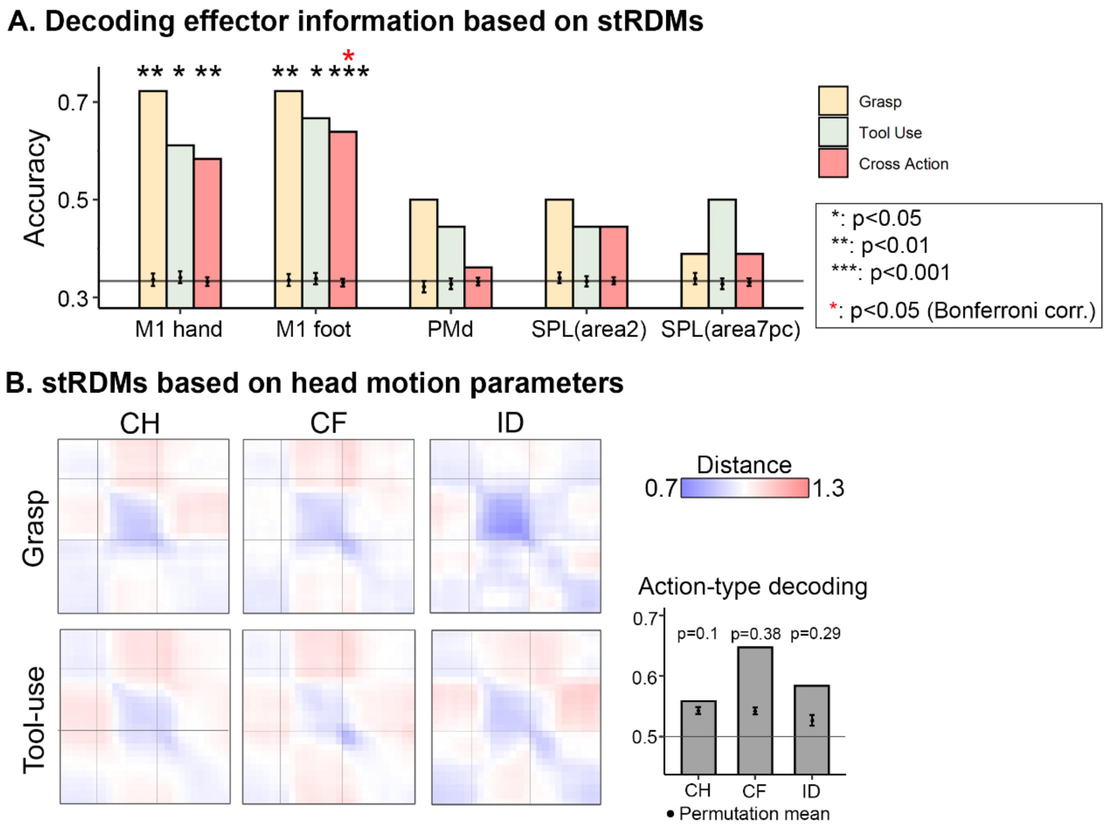


**Figure. S10. A. Decoding effector stRDMs within and across action-types in each ROI.** Effector information can be decoded in M1 areas but not association tool-use areas. B. stRDMs created using the six head motion parameters as the "response pattern" for each trial. The head motion stRDMs display high temporal similarity during action execution, a distinct pattern from the condition stRDMs shown in Fig. 6. The bar graph shows decoding of action-type based on head motion stRDMs, with the point denoting average accuracy across permutation iterations and the error bar 95% CI of permutation accuracies. P-values are calculated as the number of permutation accuracies that exceeded the actual accuracy.

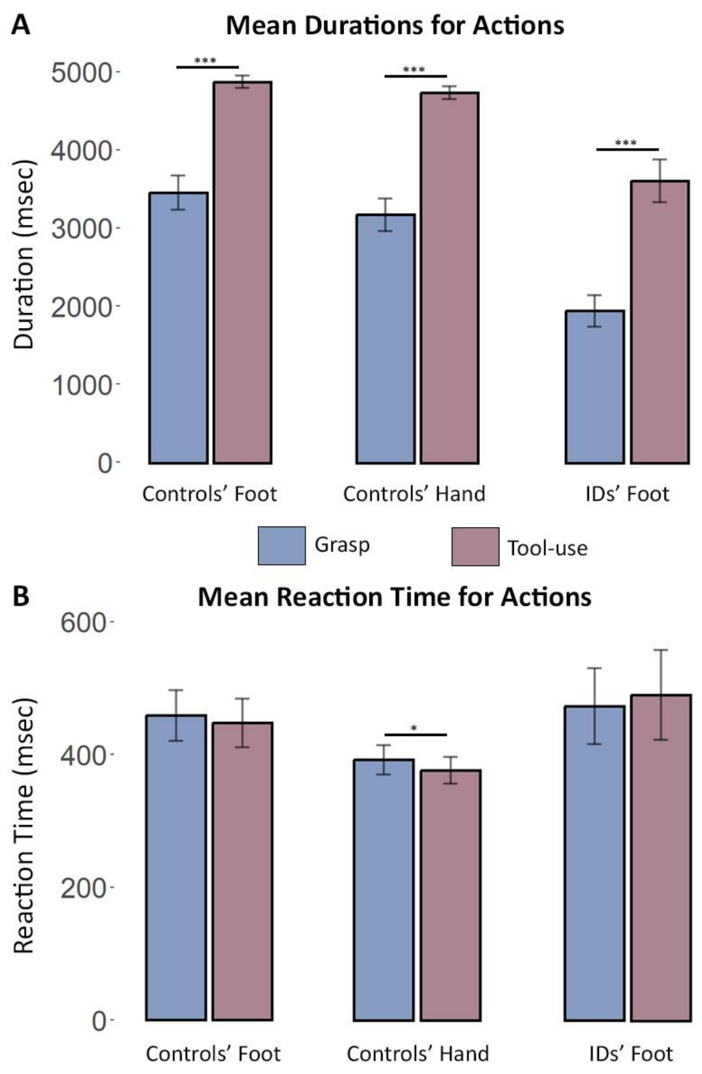


**Figure. S11. Mean Duration and Reaction Time Measurements from the Motor Behavior Recording Data.** Two accelerometers were placed on participants' executing limbs' digits (D1 and D2, corresponding to the thumb or big toe and index finger or second toes, respectively) to capture acceleration information. Two independent raters made determinations of beginning and end of movement, and their ratings were averaged. The duration (top row) was calculated by subtracting the averaged beginning of movement from the averaged end of movement. The reaction time (bottom row) was calculated by subtracting the beginning of movement from the onset cue (a beep). Measurements are captured in milliseconds. To determine if there was a difference between action-type (grasp = blue; tool-use = magenta) and executing body part/group, linear mixed effects models with fixed factors of action-type and trial and a random slope for subject were used. There was a significant difference in action duration between the two action-types, regardless of executing body part/group. There was a significant difference for reaction time between the two actions only in the controls' hand data. (* = $p < 0.05$; *** = $p < 0.001$).

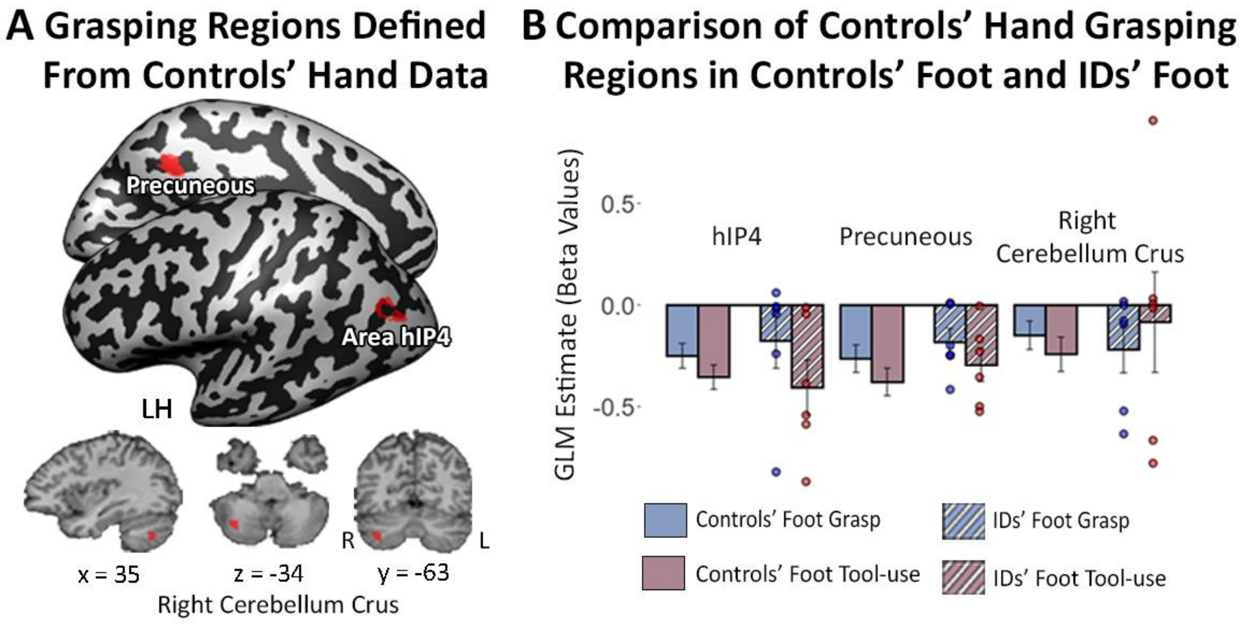


**Figure. S12. Grasping Regions as Defined by Controls' Hand Data.** These are regions that show a preference for grasping over tool-use in controls' hand data (in addition to Fig. 2A, S3; the map was thresholded at $p < 0.01$ and corrected for multiple comparisons). Bar graphs of average beta values for grasping ROIs shown in the whole-brain for controls' foot (solid bars) and IDs (striped bars). Each dot above a striped bar plot represents one individual with dysplasia. The blue color corresponds to grasp and the red color corresponds to tool-use. Information about these ROIs is presented in Table S7.

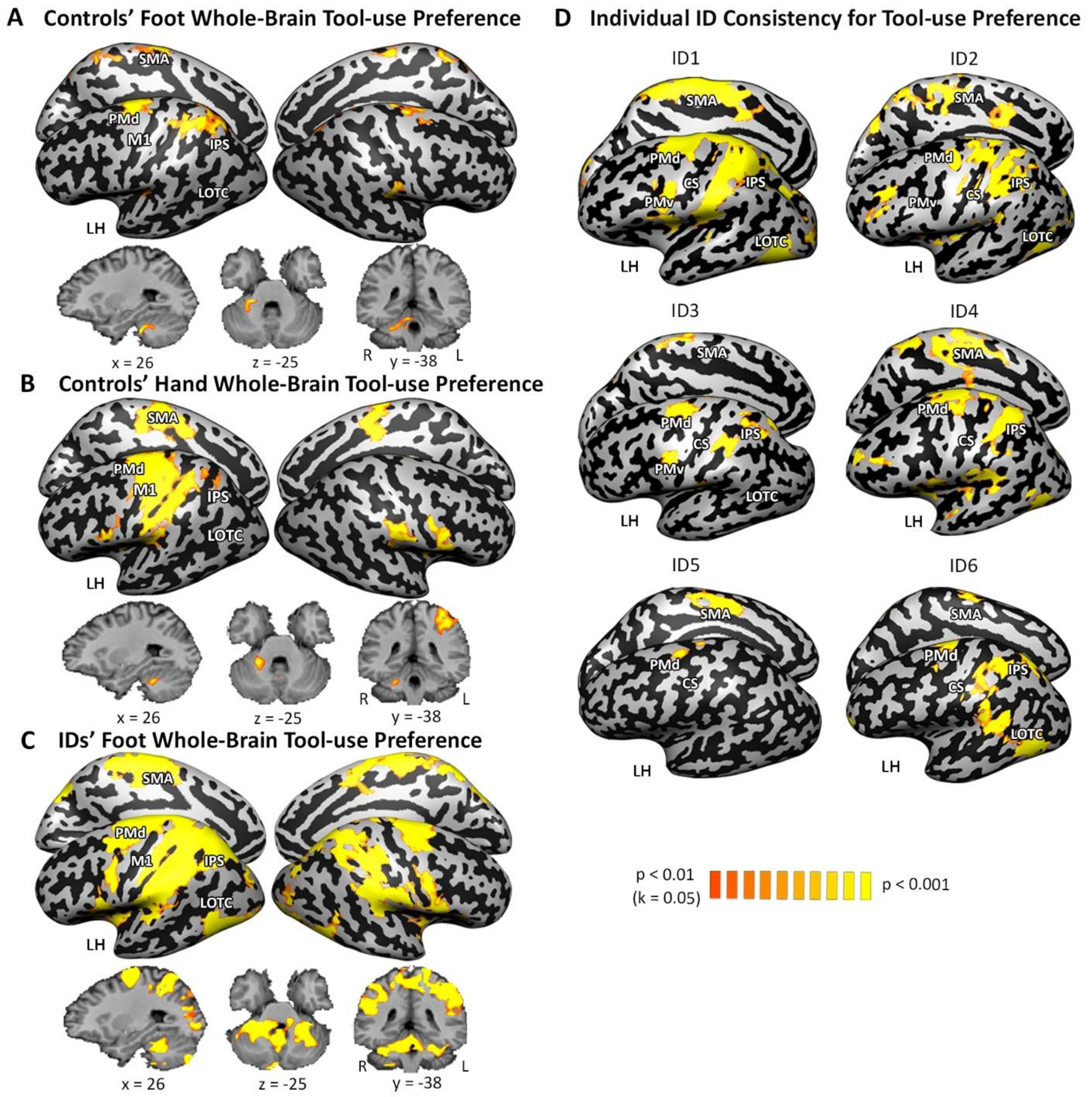


**Figure. S13. Individual Group and Individual ID Whole Brain Maps Continue to Show Whole-Brain Tool-use Preference.** Individual maps were thresholded at $p < 0.01$ and corrected for multiple comparisons at $p < 0.05$. (A,B,C) Whole-brain tool-use preference for controls' foot, controls' hand and IDs' respectively. (D) All six IDs continue to have a preference for tool-use over grasping in the PMd.

### Supplementary Tables

#### Table S1. Regions of Interest (ROIs) with a Consistent Preference for Tool-use.

| ROI name | Talairach Coordinates | | | | | | ROI Size |
|---|---|---|---|---|---|---|---|
| | X | Y | Z | Std x | Std y | Std z | Nr of Voxels |
| PMd | -26 | -18 | 62 | 2.08 | 2.14 | 2.53 | 472 |
| PMv | -53 | -4 | 38 | 1.5 | 1.91 | 2.62 | 246 |
| Area 7pc (SPL) | -30 | -58 | 52 | 2.17 | 2.65 | 2.42 | 572 |
| LOTC | -42 | -66 | -1 | 2.56 | 2.44 | 2.41 | 477 |
| Area 2 (SPL) | -38 | -42 | 50 | 1.81 | 1.69 | 2.42 | 292 |
| SMA | -5 | -13 | 54 | 1.99 | 2.32 | 2.17 | 388 |

Areas are based on the peak of the main effect of action-type from controls' hand data (thresholded at $p < 0.01$ and cluster-corrected; table for regions seen in Fig. 3). These areas were further defined as tool-use regions by looking at the controls' hand beta values from an ROI-GLM. Regions shown here had a consistent preference for tool-use as defined by ROI-GLMs for controls' foot and IDs' foot on hand tool-use areas (see Table S3 for statistics). For additional tool-use areas see Fig. S5 and Table S2.

#### Table S2. Additional Tool-use Areas

| ROI name | Talairach Coordinates | | | | | | ROI Size |
|---|---|---|---|---|---|---|---|
| | X | Y | Z | Std x | Std y | Std z | Nr of Voxels |
| Area 4p | -36 | -23 | 47 | 2.5 | 2.75 | 2.33 | 567 |
| Central Opercular Cortex | -40 | -5 | 13 | 2.53 | 1.88 | 1.95 | 359 |
| Area OP1 [S2] | -46 | -26 | 19 | 2.68 | 2.28 | 2.45 | 512 |
| Area 2 (SMG) | -49 | -27 | 38 | 2.19 | 2.55 | 2.46 | 527 |
| Right Cerebellum V | 17 | -45 | -18 | 2.74 | 2.56 | 1.93 | 506 |

Areas are based on the peak of the main effect of action-type from controls' hand data (thresholded at $p < 0.01$ and cluster-corrected; table for regions seen in Fig. S5). These areas were further defined as tool-use regions by looking at the controls' hand beta values from an ROI-GLM. For grasping regions see Fig. S12 and Table S7.

**Table S3. Significant Tool-use Region Statistics**

| Group | Area | t-value | p-value (FDR-corrected) |
|---|---|---|---|
| Controls Foot (CF, df=16) | Supplementary Motor Area (SMA) | 4.18 | $p < 0.01$ |
| | Premotor Dorsal Area (PMd) | 9.69 | $p < 0.001$ |
| | Superior Parietal Lobe (SPL Area 2) | 4.44 | $p < 0.01$ |
| | Superior Parietal Lobe (SPL Area 7pc) | 9.11 | $p < 0.001$ |
| | Lateral Occipital Temporal Cortex (LOTC) | 4.22 | $p < 0.01$ |
| | Premotor Ventral Area (PMv) | 3.57 | $p < 0.01$ |
| Individuals with Dysplasia (IDs, df=5) | Supplementary Motor Area (SMA) | 3.44 | $p < 0.05$ |
| | Premotor Dorsal Area (PMd) | 3.64 | $p < 0.05$ |
| | Superior Parietal Lobe (SPL Area 2) | 3.76 | $p < 0.05$ |
| | Superior Parietal Lobe (SPL Area 7pc) | 4.70 | $p < 0.05$ |
| | Lateral Occipital Temporal Cortex (LOTC) | 3.60 | $p < 0.05$ |
| | Premotor Ventral Area (PMv) | 3.39 | $p < 0.05$ |

Statistics from tool-use regions probed in Fig. 3B

**Table S4. Regions that decoded action-type across controls' hand and foot used in ROI MVPA**

| | Talairach Coordinates | | | | | | ROI Size |
|---|---|---|---|---|---|---|---|
| ROI name | X | Y | Z | Std x | Std y | Std z | Nr of Voxels |
| PMd | -26 | -16 | 60 | 5.91 | 5.84 | 5.76 | 3691 |
| PMv | -53 | 0 | 35 | 1.9 | 2.21 | 2.662 | 315 |
| SPL | -31 | -54 | 48 | 9.82 | 10.47 | 6.55 | 10000 |
| LOTC | -48 | -59 | 1 | 4.37 | 4.63 | 5.31 | 3028 |
| SS_3 | -53 | -24 | 39 | 3.66 | 2.12 | 2.78 | 540 |
| PFcm | -53 | -32 | 23 | 6.18 | 3.87 | 4.67 | 3027 |
| SMA | -7 | -20 | 47 | 3.31 | 3.79 | 3.46 | 1080 |
| Area hIP7 | -23 | -79 | 28 | 2.37 | 1.66 | 1.99 | 228 |
| pSTG | -53 | -42 | 16 | 2.73 | 2.12 | 3.96 | 647 |
| SS_1 | -27 | -37 | 63 | 3.45 | 4.24 | 2.52 | 826 |
| SS_2 | -50 | -18 | 46 | 2.33 | 1.61 | 2.29 | 259 |

**Table S5. Characteristics of Individuals with Dysplasia**

| Subject ID | Age | Gender | Causes of Dysplasia | Hand Prosthesis Use Experience | Upper Limb Structure |
|---|---|---|---|---|---|
| ID1 | 27 | F | Unknown | None | One residual finger attached to each shoulder |
| ID2 | 62 | M | Thalidomide | Past use of functional and cosmetic prostheses | Completely missing upper limbs bilaterally |
| ID3 | 44 | M | Unknown | Past and current occasional use of functional prostheses | Shortened right arm (± 10 cm humerus) |
| ID4 | 21 | M | Unknown | Past use of cosmetic prostheses. No use of functional protheses | One residual finger attached to the shoulder |
| ID5 | 61 | M | Thalidomide | None | Completely missing upper limbs bilaterally |
| ID6 | 38 | F | Unknown | None | Completely missing upper limbs bilaterally |

**Table S6. Identification number of IDs in current and previous studies**

| Current study | Liu et al., 2020 | Striem-Amit et al., 2017; 2018; Vannuscorps et al., 2019 (Refs.2-4) | Vannuscorps & Caramazza 2016a (5) | Vannuscorps & Caramazza 2016b (6) |
|---|---|---|---|---|
| ID1 | D1 | D1 | D5 | |
| ID2 | D2 | D2 | D2 | |
| ID3 | D3 | D4 | D3 | D5 |
| ID4 | | | | |
| ID5 | | | | |
| ID6 | | | | |

**Table S7. Grasping Areas**

| | Talairach Coordinates | | | | | | ROI Size |
|---|---|---|---|---|---|---|---|
| ROI name | X | Y | Z | Std x | Std y | Std z | Nr of Voxels |
| Area hIP4 (IPS) | -36 | -77 | 25 | 1.64 | 1.99 | 2.1 | 290 |
| Precuneous | -11 | -49 | 34 | 1.61 | 2.63 | 2.19 | 403 |
| Right Cerebellum Crus I | 35 | -63 | -34 | 2.31 | 2.08 | 2.19 | 392 |

Areas are based on the peak of the main effect of action-type from controls' hand data (thresholded at $p < 0.01$ and cluster-corrected; table for regions seen in Fig. S12). These areas were further defined as tool-use regions by looking at the controls' hand beta values from an ROI-GLM.

**Supplementary Methods**

Participants: Participant ID1 had one residual finger attached to each shoulder. ID2, ID5, and ID6 had bilateral dysplasic malformations with completely missing upper limbs on both sides (a complete absence of arm, forearm, hand and fingers). ID3 had a shortened right arm (± 10 cm humerus). Participants ID1, ID2, ID3, ID4, ID5, and ID6 apart from the congenitally missing hands, had a typically developed body.

Participant ID1 reported no history of prosthesis use. ID2 occasionally used a wood composite prosthesis with locking elbow and hooks controlled by cables attached to leg straps from 3 to 7 years old, a wood composite prosthesis with electronic elbow and three pronged hooks controlled by micro switches in shoulder harness from 7 to 11 years old and a composite prosthesis with myoelectric elbows and cosmetic hands from 11 to 15 years old. ID3 used switch-based right and left arms prostheses as a child and still uses occasionally a switch-based right arm prosthesis as an adult. ID4 used cosmetic prosthetics in the past.

All the subjects who have used prostheses report having used these prostheses mainly, if not uniquely, to pull, maintain in place or push objects but not to manipulate, and used objects for their functional use (e.g., eating with a fork) with their feet.

The IDs' profiles have been reported in previous studies (see **Table S6** for corresponding identification numbers; see **Table S5** for a summary of this information).

**References**


1. Liu, Y., Vannuscorps, G., Caramazza, A., & Striem-Amit, E. (2020). Evidence for an effector-independent action system from people born without hands. *Proceedings of the National Academy of Sciences, 117*(45), 28433-28441.
2. Striem-Amit, E., Vannuscorps, G., & Caramazza, A. (2017). Sensorimotor-independent development of hands and tools selectivity in the visual cortex. *Proceedings of the National Academy of Sciences, 114*(18), 4787-4792.
3. Striem-Amit, E., Vannuscorps, G., & Caramazza, A. (2018). Plasticity based on compensatory effector use in the association but not primary sensorimotor cortex of people born without hands. *Proceedings of the National Academy of Sciences, 115*(30), 7801-7806.
4. Vannuscorps, G., F Wurm, M., Striem-Amit, E., & Caramazza, A. (2019). Large-scale organization of the hand action observation network in individuals born without hands. *Cerebral Cortex, 29*(8), 3434-3444.
5. Vannuscorps, G., & Caramazza, A. (2016a). The origin of the biomechanical bias in apparent body movement perception. *Neuropsychologia, 89*, 281-286.
6. Vannuscorps, G., & Caramazza, A. (2016). Typical action perception and interpretation without motor simulation. *Proceedings of the National Academy of Sciences, 113*(1), 86-91.
